# Astrocyte-secreted Cathepsin D regulates hippocampal memory through extracellular matrix remodeling

**DOI:** 10.64898/2026.11.20.768232

**Authors:** Hyeyeon Kim, Seeun Cha, Minyoung Ji, Gyu Hyun Kim, Song A Shin, Seung Eun Lee, Kea Joo Lee, Sang-Won Lee, Sehyun Chae, Hyungju Park

## Abstract

Astrocytes contribute to hippocampal cognitive functions, but how astrocyte-derived factors regulate memory formation remains unclear. Here, we identify astrocyte-secreted Cathepsin D (CtsD) as a regulator of hippocampal learning and memory. Proteomic profiling of hippocampal astrocytes upon learning identified candidates predicted to be secreted, among which we focused on CtsD, whose extracellular brain functions are unknown. Neutralizing extracellular CtsD or astrocyte-selective CtsD knockdown (aCtsD-KD) impaired hippocampal long-term potentiation (LTP) and memory acquisition, suggesting an extracellular role for astrocytic CtsD. In aCtsD-KD mice, these deficits were accompanied by alterations in excitatory transmission and synaptic structures. aCtsD-KD also caused abnormal extracellular matrix (ECM) accumulation, and its partial digestion by chondroitinase ABC rescued the memory deficit, indicating that astrocytic CtsD-dependent ECM degradation contributes to hippocampal memory. Together, our findings suggest a CtsD-dependent astrocyte–matrix–synapse axis regulating hippocampal memory.

## Introduction

Astrocytes are essential components of the tripartite synapse, closely interacting with both presynaptic and postsynaptic structures throughout the central nervous system^1^. Beyond providing metabolic and structural support, astrocytes actively regulate synapse formation, elimination, and homeostasis throughout life^1–3^. In particular, astrocytes are required for the synaptic plasticity that underlies learning and memory^1^. However, it remains to be fully elucidated which molecular signals from astrocytes mediate these effects.

Among the diverse mechanisms by which astrocytes regulate synaptic functions, secreted proteins play a central role in instructing synapse development and function^1^. Previous studies have demonstrated that astrocyte-derived factors promote excitatory synaptogenesis by thrombospondins^4^ and Sparcl1^5^ or functional synapses by glypicans 4 and 6 (ref. 6) in the developing brain. In the adult hippocampus, astrocyte-derived interleukin-33 (IL-33) promotes excitatory synapse formation and is required for both homeostatic synaptic plasticity and memory formation^7^. Inhibitory synapse development is also instructed by astrocyte-secreted factors such as neurocan^8,9^. Together, diverse astrocyte-secreted proteins support the formation, maturation, and plasticity of both excitatory and inhibitory synapses. However, which astrocytic proteins are secreted in response to learning stimuli and whether such learning-induced secretory proteins are necessary for memory formation remain largely unexplored.

One structural target of astrocyte-secreted factors is the ECM, a dynamic scaffold that significantly influences synaptic structure and function^10^. Among ECM components, either chondroitin sulfate proteoglycans (CSPGs) or perineuronal nets (PNNs) are known to stabilize synaptic contacts and constrain structural plasticity and memory^11,12^. Activity-dependent ECM remodeling involves extracellular proteolysis. In the hippocampus, matrix metalloproteinase-9 (MMP-9) supports dendritic spine enlargement and LTP^13^, and a proprotein convertase–ADAMTS4/5 cascade mediates activity-dependent brevican cleavage and structural plasticity^14^. Glial cells also regulate ECM remodeling in response to neuronal activity. Microglia, for example, participate in ECM remodeling through engulfment^15^ or degradation of perisynaptic ECM through cell-surface metalloproteinase-14 (MMP-14)^16^. In contrast, the contribution of astrocyte-derived proteases to ECM regulation in the adult hippocampus remains poorly defined, although astrocytes are a major source of ECM components, including proteoglycans and tenascins^17^.

Exploring such astrocyte-derived extracellular proteins requires integrating astrocyte-targeted proteomics after learning with secretome profiling of stimulated astrocytes. To capture astrocyte-selective proteomes *in vivo*, we utilized a proximity labeling technique with TurboID^18^, which was targeted to the endoplasmic reticulum (ER) lumen of astrocytes. This approach was applied to label and isolate astrocytic proteins in the hippocampus following contextual fear conditioning. Among the learning-induced candidates predicted to be secreted, we focused on Cathepsin D (CtsD), a lysosomal aspartyl protease^19^, because neither its proteolytic activity toward ECM substrates nor the physiological function of secreted astrocytic CtsD in the brain has been characterized. Our results show that astrocytic CtsD is not only required for hippocampus-dependent memory and LTP but is also involved in ECM degradation. Partial digestion of the over-accumulated hippocampal ECM was sufficient to restore the memory deficit in astrocytic CtsD-knockdown mice. Together, these findings reveal an astrocyte–matrix–synapse axis through which astrocyte-secreted CtsD enables hippocampal memory formation.

## Results

### Hippocampal learning is associated with changes in astrocytic proteins linked to extracellular matrix and synapse organization

We first sought to profile the learning-induced astrocytic proteome of the hippocampus. To achieve selective labeling and identification of astrocytic proteins, we utilized a TurboID construct containing a KDEL sequence to direct its localization into the ER lumen (TurboID-ER) and a knock-in mouse line^18,20^ carrying a Cre-inducible *LoxP-Stop-LoxP-TurboID-ER* cassette (*LSL-TurboID-ER)*.

To test whether the TurboID-ER construct effectively labels astrocyte-derived proteins, we transduced primary mouse astrocytes with a lentivirus encoding TurboID-ER conjugated with enhanced green fluorescent protein (EGFP) under the control of the GFAP promoter (LV-Gfap-TurboID-ER-p2A-EGFP; Figure S1A). Biotinylated proteins were successfully detected in both primary astrocyte lysates and astrocyte-conditioned media (ACM; Figures S1B and S1C). We then generated an AsTER (*<u>A</u>*strocyte-*<u>s</u>*pecific *<u>T</u>*urboID-*<u>ER</u>*) mouse line by crossing the Aldh1l1-CreERT2 driver with the *LSL-TurboID-ER* line (Figure 1A). Both TurboID-ER expression and biotinylation were localized exclusively to S100β⁺ astrocytes within the hippocampus of AsTER mice (Figures 1B and S2). Furthermore, biotinylated proteins were detected in both hippocampal lysates and cerebrospinal fluid from AsTER mice following five consecutive days of intraperitoneal biotin injections (Figures S1D–S1J). Collectively, these results indicate that astrocytic expression of TurboID-ER *in vitro* and *in vivo* can label astrocyte-derived proteins.

**Figure 1.**
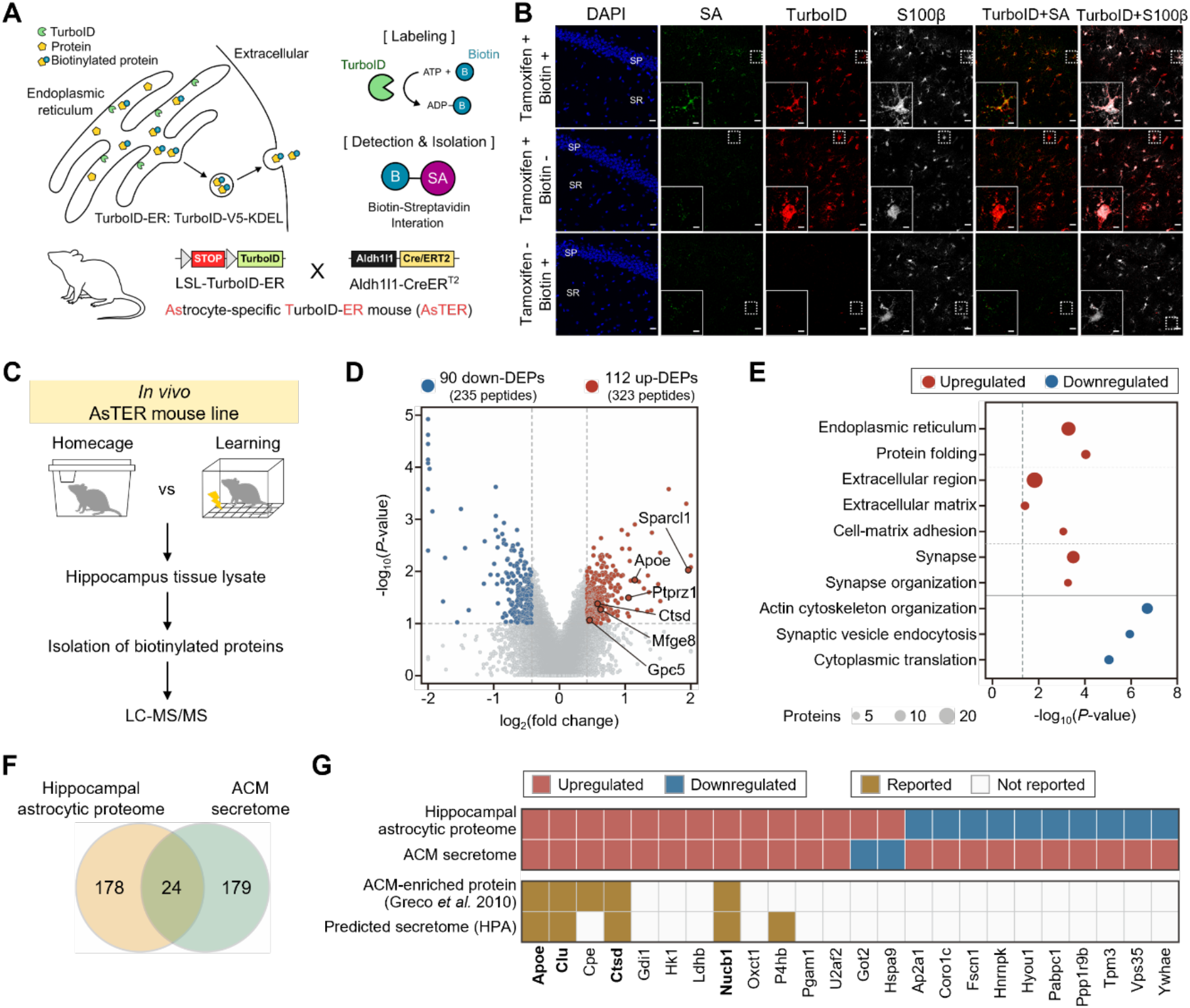
Identification of astrocytic proteins during hippocampal memory formation. **A.** Illustration of the TurboID-ER tool for astrocytic protein labeling and the AsTER mouse line. **B.** Representative confocal images of CA1 in the hippocampus showing tamoxifen- and biotin-dependent labeling of astrocytic proteins using AsTER mice. Scale bar, 20 μm; insets, 5 μm. SA, streptavidin; SP, stratum pyramidale; SR, stratum radiatum. **C.** Schematic illustration of the workflow for identifying astrocytic proteins *in vivo* using AsTER mice. **D.** Volcano plot of proteins quantified in the hippocampal astrocytic proteome of learning versus homecage AsTER mice. The x- and y-axes present the log_2_-fold change and –log_10_(*P*-value), respectively. Red and blue dots represent upregulated and downregulated proteins, respectively; gray dots represent proteins without significant differences in expression. Representative ECM-related proteins are highlighted. **E**. Representative gene ontology terms enriched in upregulated and downregulated DEPs. Circle size indicates the number of proteins. The dashed line indicates *P* = 0.05. **F.** Venn diagram showing the comparison between DEPs of the hippocampal astrocytic proteome and the ACM secretome. **G.** Heat map of the 24 overlapping proteins. Top, changes in the hippocampal astrocytic proteome (learning versus homecage) and the ACM secretome (glutamate-treated versus untreated); red, upregulated; blue, downregulated. Bottom, annotation in a published ACM-enriched proteome^24^ and the predicted secretome of the Human Protein Atlas^25^.

We next investigated learning-induced astrocytic proteomes using AsTER mice. Long-term memory formation depends on *de novo* protein synthesis^21^, and previous studies have identified an early period around the time of training and a delayed period approximately 4 h after training^22, 23^. We thus hypothesized that profiling the hippocampal astrocytic proteome 4 h after contextual fear conditioning (CFC) would capture learning-associated changes in astrocytes. AsTER mice were divided into three groups: a learning group subjected to foot shocks paired with the conditioning context, a homecage control group that remained unmanipulated, and a vehicle control (PBS) group that received PBS instead of biotin while undergoing the same training as the learning group (Figure 1C). 4 h post-training, hippocampal tissues were harvested, and biotinylated proteins were then enriched from harvested hippocampal extracts using streptavidin-coated magnetic beads (Figures S3A and S3B).

Enriched biotinylated proteins were analyzed by tandem mass tag (TMT)-based quantitative liquid chromatography-tandem mass spectrometry (LC-MS/MS), which identified 44,249 non-redundant peptides corresponding to 2,755 proteins supported by at least two unique peptides in the hippocampal astrocytic proteome (Table S1). These included 131 peptides carrying biotinylated lysine residues, validated by manual inspection of their MS/MS spectra, which directly confirm *in vivo* labeling by TurboID-ER. Gene ontology analysis showed significant enrichment for the ER and other secretory pathway compartments (Figure S3C), consistent with the localization of TurboID-ER to the ER lumen of astrocytes. Several established astrocyte-secreted proteins^24^ were among the highly abundant proteins detected (Figure S3D). These results indicate that TurboID-ER labeling in AsTER mice can capture secretory proteins of hippocampal astrocytes.

We next examined which of these astrocytic proteins responded to learning. Comparing hippocampal astrocytic proteomes from learning and control groups, we identified 202 differentially expressed proteins (DEPs), of which 112 were upregulated and 90 were downregulated (Figure 1D and Table S2). The upregulated proteins were significantly enriched for ER-related processes, including the endoplasmic reticulum and protein folding, suggesting enhanced processing of newly synthesized proteins entering the astrocytic secretory pathway upon learning. In addition to this biosynthetic signature, ECM-related processes (extracellular region, extracellular matrix, and cell-matrix adhesion) and synapse-related processes (synapse and synapse organization) were also enriched upon learning (Figure 1E). By contrast, downregulated proteins were enriched for actin cytoskeleton organization, synaptic vesicle endocytosis, and cytoplasmic translation (Figure 1E). Together, these results indicate that learning engages the astrocytic secretory pathway and selectively alters its ECM-, synapse-, and structural remodeling-related components.

To determine which of these learning-induced proteins are released from astrocytes, we performed additional proteomic analyses of ACM harvested from cultured primary mouse astrocytes, which were treated with glutamate to evoke activity-dependent protein secretion (Figure 3E). Secreted proteins were analyzed using label-free quantitative LC-MS/MS, which identified 13,591 non-redundant peptides corresponding to 1,293 proteins supported by at least two unique peptides in the ACM (Table S3). Comparing the ACM proteomes of glutamate-treated and untreated astrocytes, we identified 203 DEPs, of which 170 were upregulated and 33 were downregulated (Figure 3F and Table S4). The upregulated proteins were significantly enriched for ECM-related processes (extracellular region and extracellular matrix organization) and synapse-related processes (synapse, synaptic vesicle and neuron projection development), consistent with the ECM- and synapse-related signatures of the learning-induced hippocampal astrocytic proteome (Figure S3G).

**Figure 2.**
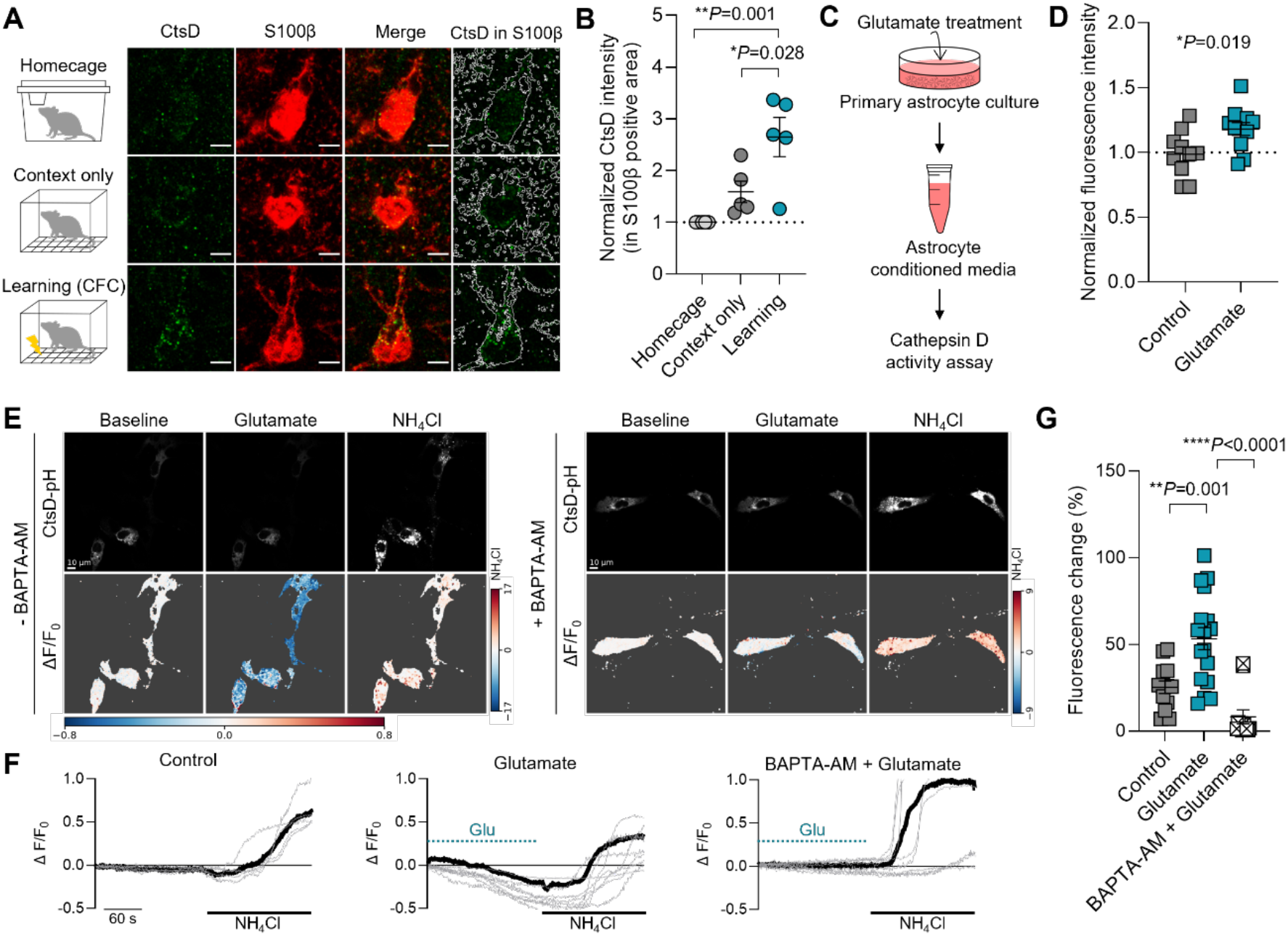
Astrocytic CtsD increases after learning and is released in an activity-dependent manner. **A**. Representative confocal images of the hippocampal CA1 region stained for CtsD (green) and astrocytes (S100β; red). White outlines indicate astrocyte areas. Scale bar, 5 μm. **B.** Quantification of CtsD fluorescence intensity within astrocyte areas. Each dot represents the mean value from one mouse. One-way ANOVA followed by Tukey’s multiple-comparisons test. Homecage, n = 5 mice; Context only, n = 5 mice; Learning, n = 5 mice. **C.** Schematic diagram of the CtsD activity assay using ACM. **D.** Quantification of the fluorescence-intensity ratio, reflecting CtsD activity, in ACM from control- and glutamate-treated astrocytes. Unpaired two-tailed Student’s *t* test. Control, n = 11 ACM samples; Glutamate, n = 10 ACM samples. **E**. Representative live-cell monitoring results acquired before (baseline) and after glutamate stimulation, as well as after NH_4_Cl treatment (NH_4_Cl). BAPTA-AM, cell-permeable Ca^2+^ chelator. Color scales for ΔF/F_0_ during baseline/glutamate stimulation and NH_4_Cl treatment are indicated at the bottom and right side of the images, respectively. **F.** Individual cellular changes in fluorescence intensity of CtsD-pH-expressing astrocytes. **G.** Quantification of changes in CtsD-pH intensity (fluorescence change %). Each dot represents one cell from three independent batches (control, n = 13; Glutamate, n = 17; BAPTA-AM + Glutamate, n = 12 cells). One-way ANOVA followed by Tukey’s multiple-comparisons test.

**Figure 3.**
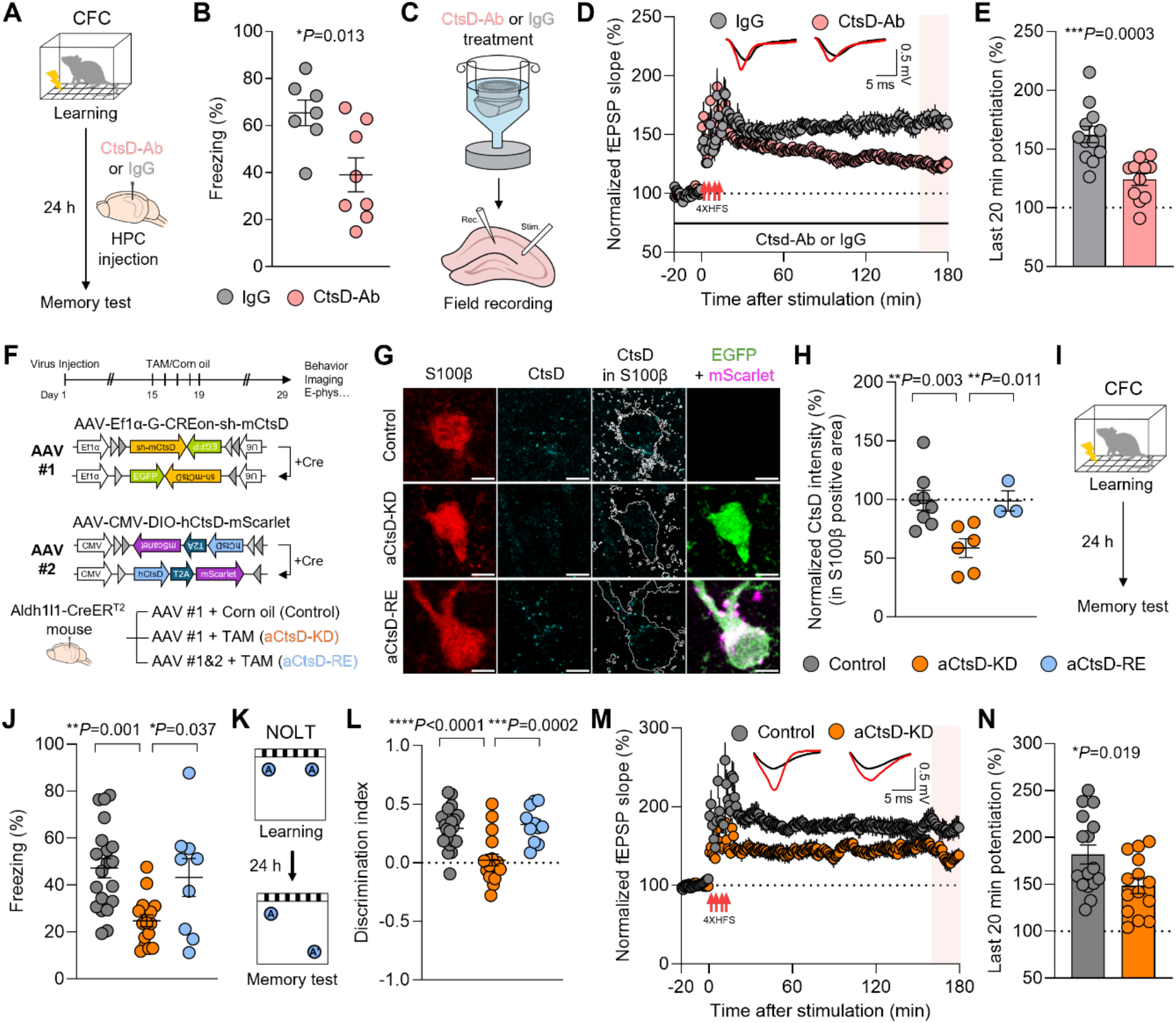
Astrocyte-secreted CtsD is required for hippocampal synaptic plasticity and memory formation. **A.** Experimental scheme for CFC and the timing of antibody injection. **B.** Percentage of time spent freezing during the CFC memory test. Unpaired two-tailed Student’s *t* test. IgG, n = 7 mice; CtsD-Ab, n = 8 mice. **C.** Schematic diagram of *ex vivo* treatment of wild-type hippocampal slices with CtsD antibody (CtsD-Ab) or control IgG. **D.** Normalized fEPSP responses over time in hippocampal slices treated with control IgG or CtsD-Ab. Red arrows indicate HFS. Representative traces obtained during baseline recording (black) and 160–180 min after HFS (red) are shown. **E.** Quantification of fEPSP responses during 160–180 min after HFS (pink box in **D**). Unpaired two-tailed Student’s *t* test. IgG, n = 12 slices from 4 mice; CtsD-Ab, n = 11 slices from 5 mice. **F.** Illustration of the astrocytic CtsD-knockdown (aCtsD-KD) and astrocytic CtsD-rescue (aCtsD-RE) mouse models. AAV #1: AAV-Ef1a-G-CREon-shCtsD; AAV #2: AAV-CMV-DIO-hCTSD-T2A-mScarlet. **G.** Representative confocal images showing CtsD fluorescence within astrocytes. White outlines indicate astrocyte areas. Scale bar, 5 μm. **H.** Quantification of CtsD fluorescence intensity within astrocyte areas. One-way ANOVA followed by Tukey’s multiple-comparisons test. Control, n = 8 mice; aCtsD-KD, n = 6 mice; aCtsD- RE, n = 3 mice. **I.** Experimental scheme for CFC. **J.** Percentage of time spent freezing during the memory test. One-way ANOVA followed by Tukey’s multiple-comparisons test. Control, n = 20 mice; aCtsD-KD, n = 16 mice; aCtsD-RE, n = 9 mice. **K.** Experimental scheme for the novel object location test (NOLT). **L.** Discrimination index of NOLT. One-way ANOVA followed by Tukey’s multiple-comparisons test. Control, n = 21 mice; aCtsD-KD, n = 17 mice; aCtsD-RE, n = 11 mice. **M.** Normalized fEPSP responses over time in hippocampal slices from Control or aCtsD-KD mice. Red arrows indicate HFS. Representative traces obtained during baseline recording (black) and 160–180 min after HFS (red) are shown. **N.** Quantification of fEPSP responses during 160–180 min after HFS (pink box in **M**). Unpaired two-tailed Student’s *t* test. Control, n = 16 slices from 5 mice; aCtsD-KD, n = 14 slices from 5 mice.

We subsequently selected proteins that were differentially expressed in both datasets (DEPs from AsTER mice and ACM), identifying 24 overlapping proteins (Figure 1F). Among these, 12 exhibited consistent directional change, all of which were upregulated in both the hippocampal astrocytic proteome and the ACM secretome, whereas the remaining 12 changed in opposite directions (Figure 1G). The consistently upregulated proteins were then filtered against a published astrocyte secretome^24^ and the predicted secretome of the Human Protein Atlas^25^. Overlapping proteins supported by both resources were retained as four astrocyte-secreted candidates (Figure 1G): Apolipoprotein E (Apoe), Clusterin (Clu), Cathepsin D (CtsD), and Nucleobindin 1 (Nucb1) (Figure 1G). Apoe and Clu are apolipoproteins best known for lipid transport^26^ and extracellular chaperone activity^27^, respectively, and both are established genetic risk factors for Alzheimer’s disease^28,29^. Nucb1, a secreted calcium-binding protein that binds amyloid precursor protein, is reduced in Alzheimer’s disease brains^30^, and inhibits amyloid-β aggregation and neurotoxicity^31^. In contrast, CtsD is the only protease among the four candidates, and its extracellular role in hippocampal function remains largely unexplored. These results support the validity of our approach for identifying functionally relevant astrocytic proteins.

### Astrocytic CtsD increases after learning and is released in an activity-dependent manner

Cathepsin family proteins are essential lysosomal proteases whose dysfunction has been implicated in the aging and pathogenesis of various neurodegenerative diseases^32^. CtsD is an aspartyl protease within this family, and its loss-of-function mutations cause neuronal ceroid lipofuscinosis (CLN10) in humans as well as severe neurodegeneration in mice^19,33,34^. Although CtsD has been studied primarily for its intracellular lysosomal functions, the extracellular role of astrocyte-derived CtsD in the brain remains unclear. We therefore investigated the contribution of astrocyte-secreted CtsD to hippocampal learning and memory.

To verify learning-induced upregulation of astrocytic CtsD (aCtsD), as suggested by our proteomic analysis (Figure 1), we evaluated CtsD protein levels using immunohistochemistry. Our analysis revealed a significant increase in CtsD expression in hippocampal astrocytes but not in neurons or microglia 4 h following learning (Figures 2A, 2B, and S4). Furthermore, aCtsD levels strongly correlated with memory performance (Figures S4E–S4G), supporting the link between aCtsD induction and hippocampal memory.

Consistent with our *in vitro* ACM proteomic findings, glutamate stimulation of cultured primary astrocytes significantly increased CtsD enzymatic activity in the ACM (Figures 2C and 2D), suggesting activity-dependent secretion of CtsD from astrocytes. To directly monitor CtsD release, we performed live-cell imaging of cultured primary astrocytes expressing CtsD fused to the pH-sensitive fluorescent protein pHluorin (CtsD-pH; Figure S5). Our results showed that glutamate stimulation decreased CtsD-pH fluorescence in astrocytes compared with bath solution alone (Figures 2E–2G). Because a stimulation-induced loss of pHluorin fluorescence has been used as a readout of the exocytic release and extracellular diffusion of pHluorin-tagged cargo^35,36^, this result suggests that glutamate stimulation triggers CtsD release from astrocytes. In addition, pretreatment of CtsD-pH-expressing astrocytes with the calcium chelator BAPTA-AM abolished the glutamate-induced decrease in CtsD-pH fluorescence (Figures 2E–2G), indicating that glutamate-evoked CtsD release requires elevated intracellular Ca^2+^.

Collectively, these results indicate that aCtsD is upregulated in hippocampal astrocytes after learning and is released in an activity-dependent manner.

### Astrocyte-secreted CtsD is required for hippocampal memory formation and long-term potentiation

Next, we asked whether CtsD is required for hippocampal synaptic plasticity and memory. To test the functional involvement of CtsD in long-term memory formation, pepstatin A, an inhibitor of aspartic proteases including CtsD, was injected into the dorsal hippocampus 4 h after CFC (Figure S6A). Mice injected with pepstatin A displayed a significant reduction in memory retrieval 24 h after conditioning (Figures S6B and S6C), indicating the importance of CtsD activity for hippocampal memory consolidation. Treatment of acute hippocampal slices with pepstatin A also impaired CA3–CA1 LTP without altering basal synaptic transmission (Figures S6D–S6G).

Because pepstatin A can also inhibit other aspartic proteases in addition to CtsD, we repeated these experiments using an anti-CtsD antibody to neutralize extracellular CtsD. Intrahippocampal injection of the CtsD antibody, but not control IgG, 4 h after CFC impaired memory retrieval at 24 h post-conditioning (Figures 3A and 3B). Likewise, LTP was not sustained in hippocampal slices treated with the CtsD antibody, whereas basal synaptic transmission remained intact (Figures 3C–3E and S7A). These results indicate that extracellular CtsD is essential for LTP and long-term memory acquisition in the hippocampus.

Finally, to determine whether hippocampal LTP and memory depend on astrocyte-derived CtsD, we generated astrocyte-selective CtsD knockdown (aCtsD-KD) mice by injecting an adeno-associated virus (AAV) encoding a Cre-dependent mouse *CtsD*-targeting shRNA expression cassette (AAV-Ef1a-G-CREon-shCtsD)^37^ bilaterally into the dorsal hippocampus of Aldh1l1-CreERT2 mice. To validate the effect of aCtsD-KD on hippocampal memory, astrocytic CtsD rescue (aCtsD-RE) mice were generated by co-injecting AAV-Ef1a-G-CREon-shCtsD with an AAV enabling Cre-dependent expression of human CTSD (AAV-CMV-DIO-hCTSD-T2A-mScarlet; Figure 3F). These viral manipulations successfully achieved significant reduction and restoration of astrocytic CtsD levels in aCtsD-KD and aCtsD-RE mice, respectively (Figures 3G, 3H, and S8).

Astrocytic CtsD reduction did not lead to altered basal locomotor activity, anxiety-like behavior, or fear responses (Figures S9A–S9H). In addition, changes in GFAP intensity and astrocytic morphology, as well as overall cytokine and chemokine profiles, were insignificant in aCtsD-KD mice compared to controls (Figure S10). These results indicate that astrocytic CtsD reduction minimally impacts overall behavioral phenotypes, astrocyte reactivity, or neuroinflammation. However, aCtsD-KD mice showed both reduced contextual fear memory (Figures 3I and 3J) and impaired object location memory 24 h after learning (Figures 3K, 3L, S9I, and S9J). Restored memory performance in both tests was observed in aCtsD-RE mice, indicating a requirement of aCtsD for hippocampal memory formation (Figures 3I–3L).

We next tested whether hippocampal LTP requires astrocytic CtsD. LTP was significantly impaired in hippocampal slices from aCtsD-KD mice (Figures 3M and 3N). In contrast to slices treated acutely with pepstatin A or the CtsD antibody (Figures S6F and S7A), basal synaptic strength was also significantly reduced in aCtsD-KD slices (Figure S7B). This distinction likely reflects the chronic downregulation of CtsD in aCtsD-KD mice, which depletes both extracellular and intracellular astrocytic CtsD pools, possibly leading to broader synaptic deficits than acute extracellular blockade.

Together, these results demonstrate that astrocyte-derived extracellular CtsD is necessary for long-term synaptic plasticity and memory formation in the adult hippocampus.

### Astrocytic CtsD maintains the structural and functional integrity of hippocampal excitatory synapses

We next investigated whether basal neurotransmission is also affected by aCtsD. First, we performed whole-cell patch-clamp recordings from CA1 pyramidal neurons in hippocampal slices treated with a CtsD antibody to monitor the spontaneous excitatory and inhibitory postsynaptic currents (sEPSCs and sIPSCs, respectively). Inhibiting extracellular CtsD significantly increased the frequency of sEPSCs without affecting amplitude or the frequency and amplitude of sIPSCs (Figure S11). aCtsD-KD produced a significant increase in sEPSC frequency with no change in sEPSC amplitude, in line with the effects of extracellular CtsD blockade. However, aCtsD-KD decreased both the frequency and amplitude of sIPSCs (Figures 4A–4C). No detectable alterations were observed in the overall intrinsic membrane properties of the recorded CA1 neurons (Supplementary Table 5).

**Figure 4.**
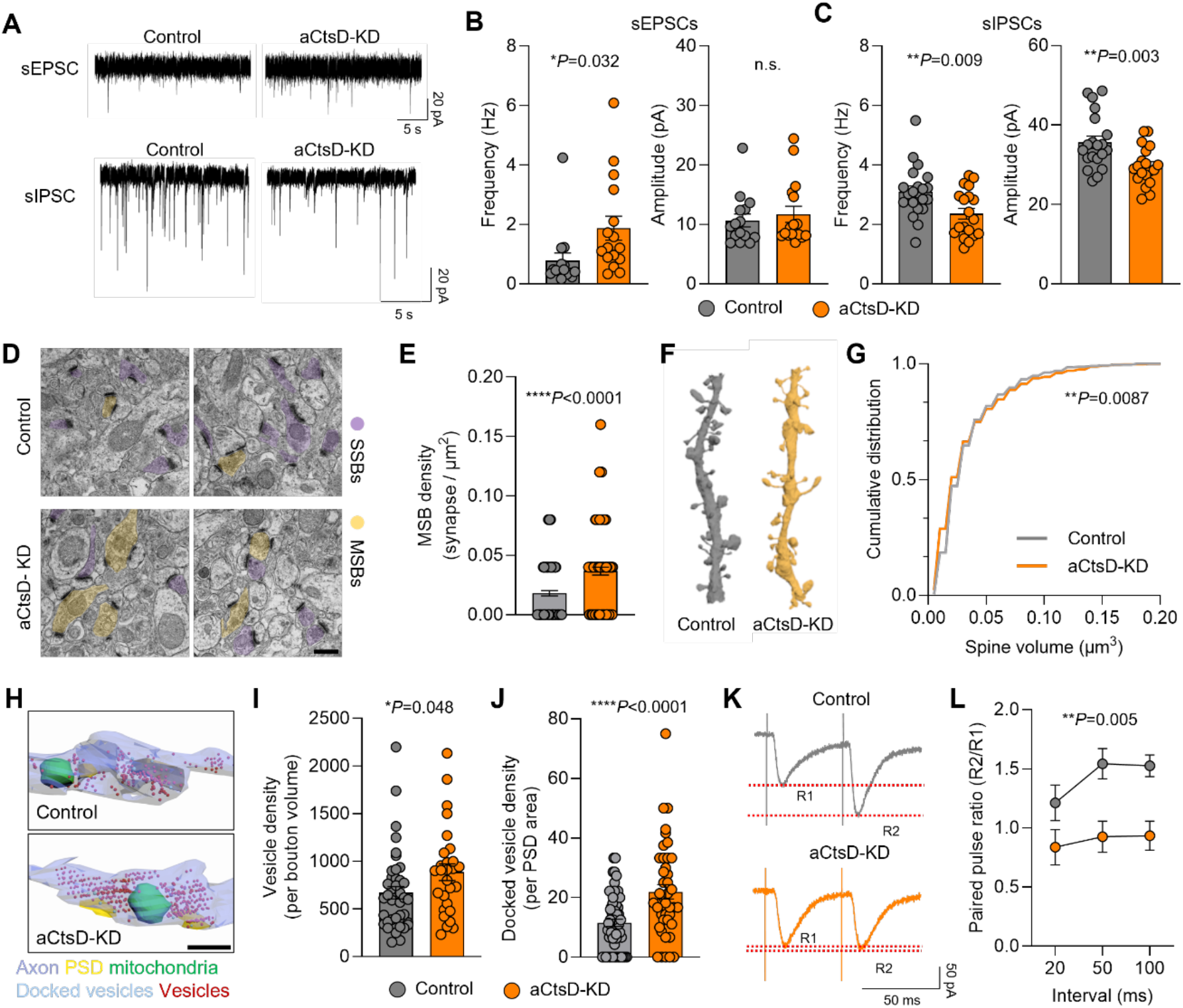
Astrocytic CtsD regulates the functional and structural integrity of hippocampal synapses. **A.** Representative traces of sEPSCs (top) and sIPSCs (bottom) recorded from hippocampal CA1 neurons in Control and aCtsD-KD mice. **B, C.** Quantification of the frequency and amplitude of sEPSCs (**B**) and sIPSCs (**C**). Unpaired two-tailed Student’s *t* test. Control, n = 15 cells from 4 mice; aCtsD-KD, n = 16 cells from 4 mice (**B**). Control, n = 20 cells from 4 mice; aCtsD-KD, n = 19 cells from 4 mice (**C**). **D.** Representative transmission electron microscopy images showing single-synaptic boutons (SSBs; purple) and multi-synaptic boutons (MSBs; yellow). Scale bar, 0.5 μm. **E.** Quantification of MSB density. Unpaired two-tailed Student’s *t* test. Control, n = 120 images from 4 mice; aCtsD-KD, n = 120 images from 4 mice. **F.** Representative three-dimensional (3D) reconstructions of dendritic spines. **G.** Cumulative distribution of spine volume (μm^3^). Kolmogorov-Smirnov test. Control, n = 539 spines from 3 mice; aCtsD-KD, n = 496 spines from 3 mice. **H.** Representative 3D reconstructions of excitatory presynaptic boutons from Control and aCtsD-KD mice, showing the spatial organization of synaptic vesicles (red), docked vesicles (light blue), postsynaptic densities (PSDs; yellow), and mitochondria (green). Scale bar, 1 μm. **I.** Quantification of total synaptic vesicle density per bouton volume. Unpaired two-tailed Student’s *t* test. Control, n = 41 boutons from 3 mice; aCtsD-KD, n = 28 boutons from 3 mice. **J.** Quantification of docked vesicles per PSD area. Unpaired two-tailed Student’s *t* test. Control, n = 59 PSDs from 3 mice; aCtsD-KD, n = 42 PSDs from 3 mice. **K.** Representative traces of paired-pulse responses recorded from hippocampal CA1 neurons in Control and aCtsD-KD mice. R1 and R2 indicate the amplitudes of the first and second evoked responses, respectively. **L.** Quantification of the paired-pulse ratio (R2/R1) at interpulse intervals of 20, 50, and 100 ms. Two-way repeated-measures ANOVA with Tukey’s multiple-comparisons test. Control, n = 13 cells from 4 mice; aCtsD-KD, n = 18 cells from 4 mice.

Despite the reduction in sIPSC frequency and amplitude, we detected no differences in the density of putative inhibitory synaptic contacts or in the number of parvalbumin-positive and somatostatin-positive interneurons between control and aCtsD-KD mice (Figures S12A and S12C–S12E). Thus, the reduced inhibitory transmission in aCtsD-KD mice is unlikely to result from a loss of inhibitory synapses or interneurons. Moreover, because acute blockade of extracellular CtsD did not alter sIPSCs (Figure S11), this inhibitory synapse phenotype may arise from the chronic loss of astrocytic CtsD, which removes both its intracellular and secreted pools, rather than from the acute loss of extracellular CtsD activity.

Because both extracellular CtsD blockade and aCtsD-KD increased sEPSC frequency, we next asked which structural features of excitatory synapses depend on aCtsD. To address this question, we performed transmission electron microscopy (TEM). Despite no change in the density of putative excitatory synaptic contacts by aCtsD-KD (Figure S12B), putative excitatory multi-synaptic boutons (MSBs; individual presynaptic terminals contacting multiple postsynaptic spines) occurred more frequently in the aCtsD-KD hippocampus (Figures 4D and 4E). This suggests that aCtsD limits aberrant structural changes in excitatory connectivity, such as MSB formation, without altering overall synapse density. To further characterize the fine ultrastructural changes occurring at excitatory pre- and postsynaptic sites, we performed automated tape-collecting ultramicrotome scanning electron microscopy (ATUM-SEM) along with 3D image reconstruction (3D-EM; Figures S13A–S13C). This analysis revealed that the spine volume of CA1 pyramidal neurons is significantly reduced in the aCtsD-KD hippocampus (Figures 4F and 4G). Corroborating this finding, TEM analysis revealed shorter postsynaptic density (PSD) lengths in aCtsD-KD mice compared to controls (Figures S13D and S13E).

Beyond postsynaptic structural remodeling, ATUM-SEM and 3D-EM analyses also revealed structural alterations in presynaptic terminals in the aCtsD-KD hippocampus. Specifically, aCtsD-KD mice exhibited significant increases in the densities of both total presynaptic vesicles and docked vesicles (Figures 4H–4J and S13B), likely contributing to enhanced presynaptic neurotransmitter release. Supporting this idea, the paired-pulse ratio at CA3–CA1 synapses was significantly reduced in the aCtsD-KD hippocampus compared to control mice (Figures 4K and 4L). Consequently, aCtsD depletion appears to induce distinct, opposing adaptations across the excitatory pre- and postsynaptic sides: reduced postsynaptic dimensions accompanied by enhanced presynaptic release. The latter mechanism may contribute to the increase in sEPSC frequency (Figures 4A and 4B) despite no change in overall synapse density (Figure S12B). These findings collectively demonstrate that aCtsD is crucial for preserving the structural and functional integrity of excitatory synapses.

### Astrocyte-secreted CtsD promotes hippocampal extracellular matrix turnover

We next explored how aCtsD regulates synaptic function and structure in the adult hippocampus through extracellular mechanisms. Studies have demonstrated that ECM remodeling contributes to activity-dependent synapse organization and plasticity^13,14,38^. Based on previous evidence implicating CtsD in ECM remodeling in peripheral tissues^39,40^, we hypothesized that extracellular aCtsD potentially controls ECM remodeling in the hippocampus.

To test this hypothesis, we evaluated *Wisteria floribunda* agglutinin (WFA)-labeled ECM structures in the hippocampus of control and aCtsD-KD mice (Figures 5A–5C). Both the area and signal intensity of WFA were significantly increased in the aCtsD-KD hippocampus (Figures 5B and 5C). Given that CtsD is a key lysosomal enzyme^19^, an intracellular reduction in CtsD by aCtsD-KD might alter lysosomal functions, thereby impairing cellular uptake and degradation of ECM. However, our results showed that intracellular depletion of aCtsD minimally affects the ECM level, because astrocyte morphology, lysosomal density, and astrocytic ECM engulfment remained unchanged in the aCtsD-KD hippocampus (Figures S14A–S14D). Furthermore, aCtsD-KD did not affect microglial morphology, microglial lysosomal density, and lysosomal inclusion of ECM components in microglia (Figures S14E–S14). Collectively, these results suggest a significant role of extracellular CtsD in ECM degradation. Supporting this idea, intrahippocampal delivery of a neutralizing CtsD antibody, but not control IgG, reproduced the over-accumulation of hippocampal ECM (Figures 5D–5F).

**Figure 5.**
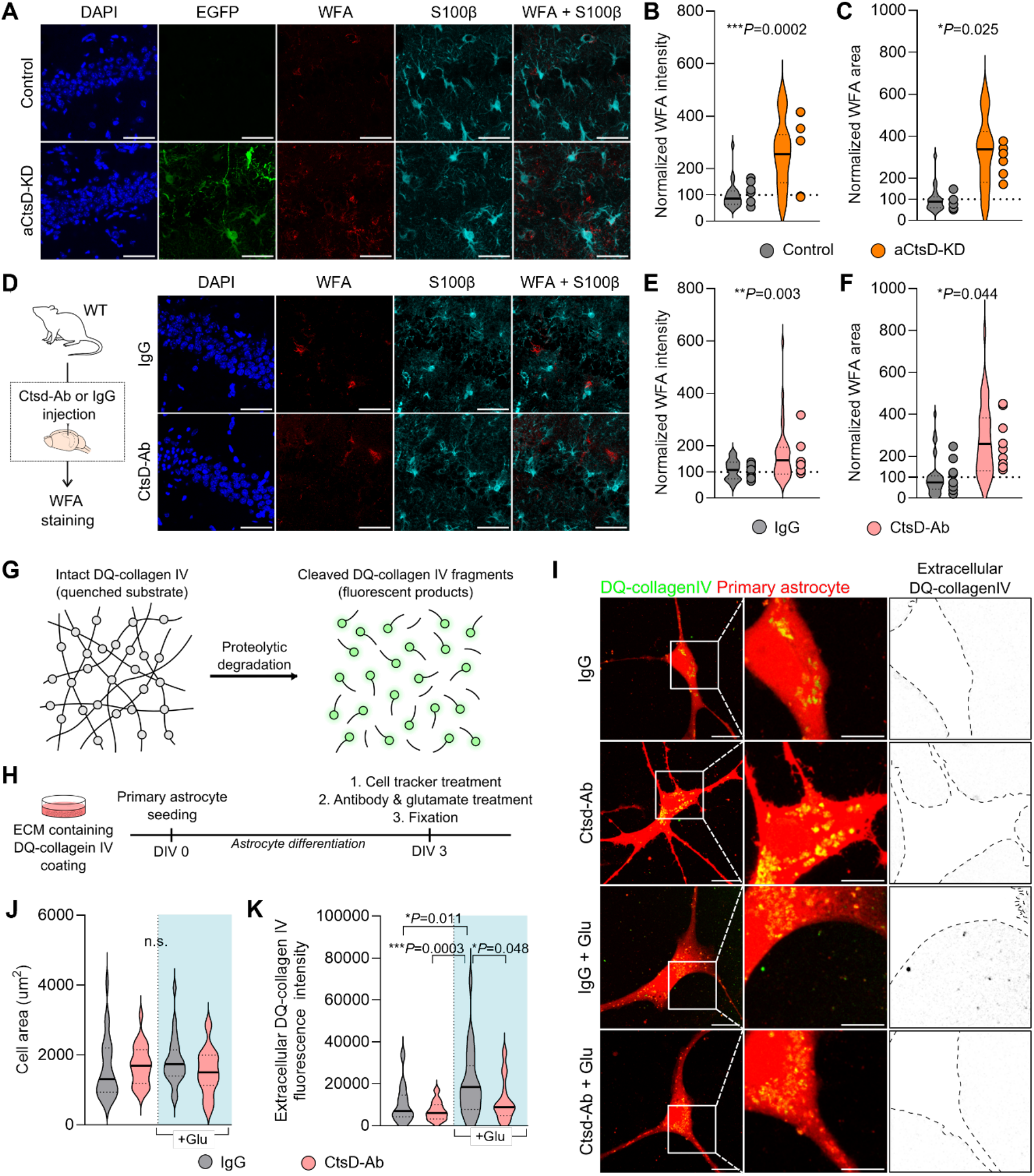
Astrocyte-secreted CtsD degrades extracellular matrix. **A.** Representative confocal images of the hippocampal CA1 region stained for DAPI (blue), EGFP (green), WFA (red), and S100β (cyan) in Control and aCtsD-KD mice. Scale bar, 50 μm. **B, C.** Quantification of the WFA fluorescence intensity ratio (**B**) and WFA-positive area ratio (**C**) in Control and aCtsD-KD mice. Unpaired two-tailed Student’s *t* test. Control, n = 6 mice; aCtsD-KD, n = 6 mice. **D.** Experimental scheme (left) for intrahippocampal injection of CtsD antibody (CtsD-Ab) or control IgG in wild-type (WT) mice, followed by WFA staining. Representative confocal images (right) of the hippocampal CA1 region stained for DAPI (blue), WFA (red), and S100β (cyan). Scale bar, 50 μm. **E, F.** Quantification of the WFA fluorescence intensity ratio (**E**) and WFA-positive area ratio (**F**) in IgG- and CtsD-Ab–injected WT mice. Unpaired two-tailed Student’s *t* test. IgG, n = 10 mice; CtsD-Ab, n = 10 mice. **G.** Schematic illustration of the DQ-collagen IV proteolysis assay. **H.** Experimental timeline of the DQ-collagen IV proteolysis assay in primary astrocytes. **I.** Representative images showing extracellular DQ-collagen IV degradation in primary astrocytes treated with IgG, CtsD-Ab, IgG + glutamate (Glu), or CtsD-Ab + Glu. Scale bars, 20 μm; insets, 10 μm. **J, K.** Quantification of cell area (**J**) and extracellular DQ-collagen IV integrated fluorescence intensity (**K**). One-way ANOVA followed by Tukey’s multiple-comparisons test. IgG, n = 24 cells; CtsD-Ab, n = 24 cells; IgG + Glu, n = 23 cells; CtsD-Ab + Glu, n = 24 cells. All cells from three different batches.

Whether astrocyte-derived extracellular CtsD contributes to ECM degradation in the near-neutral extracellular conditions remained unclear. To address this question, we used a cell-based ECM degradation assay. The C8D1a mouse astrocyte cell line was cultured on substrate-bound DQ-collagen IV, a heavily fluorescein-labeled type IV collagen matrix. Proteolytic cleavage of this self-quenched substrate restores fluorescence, providing a direct fluorescent readout of ECM breakdown^41^ (Figure 5G). Overexpression of human CTSD (hCTSD) in the C8D1a cell line significantly increased DQ-collagen IV fluorescence, and this substrate degradation was blocked by pepstatin A (Figure S15). Such pepstatin A-sensitive degradation of the extracellular collagen IV matrix, observed in standard culture medium at near-neutral pH, raises the possibility that extracellular CtsD contributes to ECM degradation.

We then assessed whether endogenous CtsD released from astrocytes can mediate ECM breakdown. Primary mouse astrocytes were plated on substrate-bound DQ-collagen IV and treated with 100 μM glutamate to evoke CtsD release (Figure 5H). Our results showed that glutamate stimulation of primary astrocytes significantly enhanced DQ-collagen IV degradation, and this activity-dependent ECM breakdown was inhibited by co-application of a CtsD-neutralizing antibody (Figures 5I–5K). Taken together, these results indicate that astrocyte-secreted CtsD participates in activity-dependent ECM breakdown by astrocytes. Whether extracellular CtsD is directly or indirectly implicated in ECM cleavage remains to be explored.

### Normalizing ECM accumulation restores memory in aCtsD-KD mice

Given that aCtsD depletion both disrupted ECM turnover and impaired memory, excessive ECM accumulation appears to mediate this memory impairment. Therefore, we next asked whether normalizing ECM levels with CtsD-independent mechanisms could ameliorate the memory impairment in aCtsD-KD mice. Over-accumulation of the hippocampal ECM in aCtsD-KD mice was inhibited by injecting chondroitinase ABC (ChABC).

To determine the optimal ChABC dose for normalizing ECM levels without eliminating the matrix, we first performed a dose titration in wild-type mice by injecting PBS into the left hippocampus and varying concentrations of ChABC into the right hippocampus. ChABC treatment led to a dose-dependent reduction in hippocampal WFA intensity compared to the contralateral PBS control, and a dose of 0.05 mU ChABC significantly attenuated the WFA signal while preserving baseline matrix level (Figures 6A and 6B). We therefore selected 0.05 mU for the subsequent functional rescue experiments.

**Figure 6.**
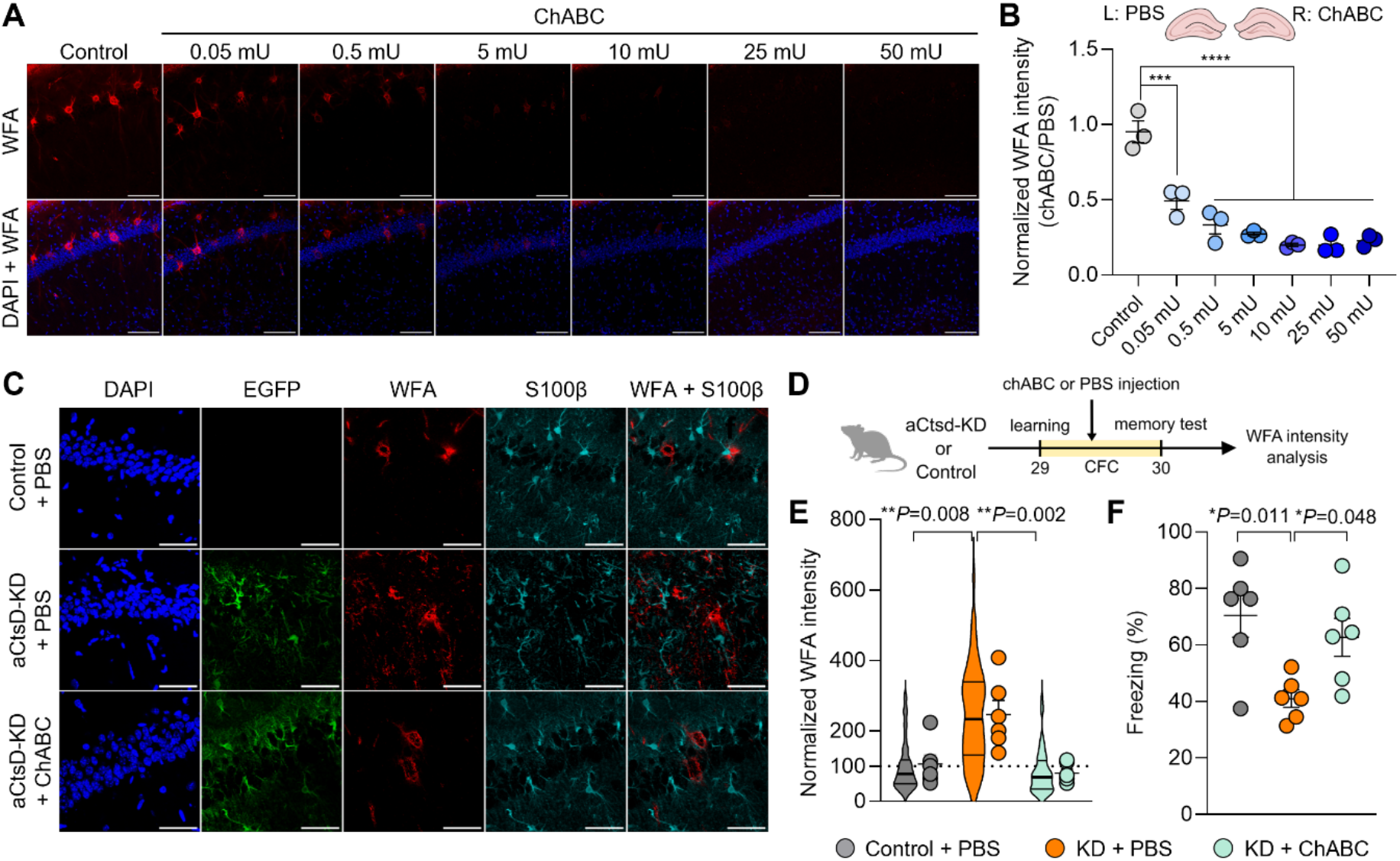
ChABC-mediated ECM normalization rescues memory deficits in aCtsD-KD mice. **A.** Representative confocal images of WFA (red) staining in the hippocampus following intrahippocampal injection of ChABC. Scale bar, 100 μm. **B.** Quantification of the WFA intensity ratio in the hippocampus following unilateral injection of phosphate-buffered saline (PBS; left hemisphere) and ChABC (right hemisphere). Each dot represents an individual mouse (n = 3 mice per concentration). One-way ANOVA followed by Tukey’s multiple-comparisons test. **C.** Representative confocal images of the hippocampal CA1 region stained for DAPI (blue), EGFP (green), WFA (red), and S100β (cyan) in Control mice injected with PBS, aCtsD-KD mice injected with PBS, and aCtsD-KD mice injected with ChABC. Scale bar, 50 μm. **D.** Experimental timeline of ChABC or PBS injection before the memory test, followed by WFA intensity analysis. **E.** Quantification of the WFA fluorescence intensity ratio. One-way ANOVA followed by Tukey’s multiple-comparisons test. Control + PBS, n = 6 mice (56 images); aCtsD-KD + PBS, n = 6 mice (54 images); aCtsD-KD + ChABC, n = 6 mice (55 images). **F.** Percentage of time spent freezing during the memory test. One-way ANOVA followed by Tukey’s multiple-comparisons test. Control + PBS, n = 6 mice; aCtsD-KD + PBS, n = 6 mice; aCtsD-KD + ChABC, n = 6 mice.

We then administered either ChABC (0.05 mU) or PBS into the hippocampus of aCtsD-KD mice 4 h after CFC. Memory retrieval was evaluated 24 h post-conditioning of CFC, followed by histological quantification of hippocampal WFA intensity (Figures 6C–6F). Intrahippocampal ChABC infusion successfully normalized WFA intensity and restored contextual fear memory in aCtsD-KD mice to control levels. In contrast, PBS-treated aCtsD-KD mice exhibited persistent ECM accumulation and pronounced memory deficits. These findings indicate that restoring ECM homeostasis is sufficient to reverse the memory impairment in aCtsD-KD mice, supporting our hypothesis that regulated ECM remodeling by astrocyte-derived CtsD is required for hippocampal memory.

## Discussion

In this study, we identified astrocyte-secreted CtsD as a novel, activity-regulated factor required for the synaptic plasticity that underlies long-term memory formation in the hippocampus. By combining *in vivo* astrocyte-specific proteomic and *in vitro* secretome analyses, we characterized the learning-induced astrocytic proteome and found that astrocytes upregulate and secrete CtsD (Figures 1, 2, S3, and S4). Our data also implicate this astrocyte-derived extracellular CtsD in the proteolytic regulation of the ECM (Figure 5). Moreover, hippocampal memory formation depended on aCtsD-mediated ECM degradation (Figures 3B, 3J, 3L, and 6). Together, these results suggest that learning-induced aCtsD supports hippocampal synaptic plasticity and memory formation through ECM degradation.

Although CtsD is best known as a lysosomal protease, the role of astrocyte-derived extracellular CtsD in learning and memory has remained largely unexplored. In non-brain tissues, extracellular CtsD has been shown to contribute to tissue remodeling and tumor-associated matrix regulation^40,42^. Because either aCtsD-KD or neutralization of extracellular CtsD caused ECM accumulation (Figures 5A–5F), we reasoned that astrocyte-derived extracellular CtsD contributes to ECM regulation in the adult hippocampus. In line with this idea, our cell-based ECM degradation assay indicated that astrocyte-derived CtsD contributes to the breakdown of a collagen IV matrix (Figures 5G–5J and S15. Whether CtsD cleaves other ECM components directly or indirectly remains to be determined.

Two non-exclusive models may explain mechanisms underlying CtsD regulation of ECM turnover. First, CtsD may directly cleave ECM components. Previous reports demonstrated that CtsD substrates include SPARC (osteonectin/BM-40)^40^, fibronectin^43^, and aggrecan^44^. Second, CtsD may act indirectly by activating extracellular protease networks. At acidic pH, CtsD can activate secreted procathepsin B^45^ and cleave cystatin C, the potent endogenous inhibitor of cysteine cathepsins, thereby augmenting extracellular cysteine cathepsin activity^46^. Moreover, extracellular CtsD may enhance extracellular plasminogen activator activity by degrading plasminogen activator inhibitor-1^47^. Future studies should delineate the precise molecular cascade underlying CtsD-induced ECM remodeling.

An important remaining question is how secreted CtsD regulates ECM turnover in the near-neutral extracellular environment of the hippocampus. Although CtsD is most active under acidic conditions, secreted CtsD can mediate limited proteolysis at physiological pH^35,48^. Moreover, intense neuronal activity can induce transient local extracellular acidification at active synapses or near astrocytic processes^49,50^, potentially influencing the catalytic functions of pH-sensitive proteins such as CtsD.

Our study does not exclude the possibility that intracellular CtsD in astrocytes also contributes to regulating synaptic function. This notion is supported by the finding that the electrophysiological consequences of chronic aCtsD-KD differed from those of acute extracellular CtsD blockade (Figures 3 and S7). Specifically, aCtsD-KD, which chronically depletes both intracellular and extracellular CtsD pools, resulted in broad synaptic deficits, including altered basal synaptic transmission and LTP, whereas selective acute blockade of extracellular CtsD impaired only LTP (Figures 3 and S7). These broader deficits in aCtsD-KD mice suggest distinct contributions of intracellular and extracellular astrocytic CtsD to synaptic function. Alternatively, the loss of lysosomal CtsD in astrocytes may secondarily alter the levels of other astrocyte-derived synaptogenic or ECM-related factors, such as SPARCL1/hevin, thrombospondins, glypicans, and neurocan^1,9^, which might contribute cell-autonomously to the synaptic phenotypes in aCtsD-KD mice.

The adult brain ECM has been widely regarded as a structural scaffold that stabilizes mature synapses^38^. However, excessive accumulation of ECM and PNN has been proposed to constrain synaptic plasticity by restricting the structural and molecular remodeling required for synaptogenesis and remodeling^51,52^. Our findings underscore the importance of dynamic ECM turnover in supporting synaptic plasticity in the mature brain, as inhibiting extracellular CtsD or knocking down astrocytic CtsD led to matrix accumulation in the hippocampus (Figure 5), alongside impairments in hippocampal LTP and memory formation (Figure 3). Crucially, partial enzymatic digestion of the over-accumulated matrix was sufficient for restoring contextual fear memory in aCtsD-KD mice (Figure 6). Together, these findings support a causal contribution of aCtsD-mediated ECM turnover to memory formation.

Our results show that aCtsD depletion altered synaptic organization in addition to ECM accumulation, suggesting that aCtsD-dependent ECM reorganization is involved in regulating synapses. In the aCtsD-KD hippocampus, excitatory synapses displayed an abnormal increase in MSB without a change in total synapse density (Figures 4D, 4E, and S12), as well as smaller postsynaptic structures, elevated presynaptic vesicle counts, and an increased release probability (Figures 4F–4L and S13). Given that structural remodeling of postsynaptic spines depends on ECM integrity^38,53,54^, the observed increase in MSB and presynaptic release strength may reflect a homeostatic response to reduced postsynaptic efficacy. Alternatively, excessive ECM accumulation may suppress integrin signaling, maintaining presynaptic terminals in a high-release-probability state^53,54^ and inducing compensatory downsizing of postsynaptic sites. As a result, the combination of ECM over-accumulation and subsequent homeostatic adaptation of excitatory synapses likely drives the synaptic plasticity deficits (Figures 3D, 3E, 3M, and 3N). Overall, these findings point to a role for astrocyte-derived CtsD in regulating ECM turnover to preserve excitatory synaptic connectivity.

Previous work has shown that microglial engulfment and MMP-14 are involved in ECM turnover and synapse dynamics in the adult and developing circuits, respectively^15,16^, suggesting the importance of microglial mechanisms for ECM regulation. On the other hand, our study reveals astrocytic ECM regulation via CtsD-dependent extracellular proteolysis and its contribution to long-term memory formation in the adult hippocampus. These findings collectively suggest that the mechanisms underlying matrix stabilization and turnover on synaptic plasticity vary depending on circuit maturation state. On the other hand, given microglia-astrocyte crosstalk instructing synapse and ECM remodeling^55,56^, hippocampal ECM integrity may be governed by complex glial mechanisms ranging from phagocytic clearance to extracellular proteolysis. Whether astrocytic CtsD-dependent remodeling interacts with microglial mechanisms remains to be determined.

The astrocyte–matrix–synapse axis identified here may also be relevant to neurodegenerative disorders, such as Alzheimer’s disease (AD)^57^, in which lysosomal degradation and synaptic maintenance are impaired. Previous studies on brain CtsD have mainly focused on its intracellular roles in proteostasis^33,34^. Aberrant ECM and PNN accumulation are also linked to AD pathogenesis^51,58^ and may contribute to synaptic dysfunction. Our results raise the possibility that neurodegeneration linked to CtsD deficiency stems not only from intracellular lysosomal failure but also from loss of its extracellular, matrix-remodeling function. Similarly, excessive PNN and CSPG deposition accompanied by reduced synaptic plasticity occurs in the aging brain^51,59^, suggesting that an age-related reduction in astrocytic CtsD secretion may contribute to cognitive decline. Further studies are needed to examine whether alterations in astrocytic CtsD secretion or extracellular activity drive pathological ECM remodeling in aging and neurodegeneration.

This study has several limitations. First, because the direct neuronal and extracellular substrates of CtsD in the brain remain unknown, the detailed molecular mechanisms underlying aCtsD-mediated matrix processing are yet to be defined. Specifically, it is unclear whether CtsD acts directly or indirectly, whether the pro- or mature form is active, and what extracellular pH microenvironment is required. Second, our analyses were performed at a single time point following CtsD manipulation, preventing us from resolving the precise temporal sequence by which aCtsD-mediated ECM remodeling affects pre-versus postsynapses. Third, due to the partial deficiency of aCtsD by shRNA-mediated knockdown, the phenotypes observed in aCtsD-KD mice may underestimate the full contribution of astrocytic CtsD.

In summary, our study identifies astrocyte-secreted CtsD as a novel, activity-regulated factor required for hippocampal memory formation. By promoting ECM degradation, astrocytic CtsD maintains dynamic ECM turnover, thereby facilitating structural and functional synaptic plasticity. These findings uncover an astrocyte–matrix–synapse axis through which an astrocyte-derived extracellular protease supports memory formation in the adult hippocampus.

## Acknowledgments

This research was supported by grants from the KBRI Basic Research Program (26-BR-01-02 and 26-BR-01-03) and by the K-Brain Project of the National Research Foundation of Korea (NRF; RS-2023-00265524) funded by the Ministry of Science and ICT. H.K. was supported by the Basic Science Research Program through the NRF funded by the Ministry of Education (RS-2026-25560987). S.-W.L. was supported by the grants funded by the Ministry of Education (RS-2022-NR068428) and the Ministry of Science and ICT (RS-2025-16652968), as well as the National Cancer Center funded by the Ministry of Health and Welfare (RS-2024-00441759). S.Chae was supported by the Global-Learning and Academic Research Institution for Master’s and PhD Students and Postdoctoral Researchers (G-LAMP) Program of the NRF funded by the Ministry of Education (RS-2023-00301850) and by the Basic Science Research Program through the NRF funded by the Ministry of Science and ICT (RS-2026-25494803).

## Author contributions

H.K. and H.P. designed the project. H.K. and S.A.S. performed virus injections and behavioral tests. H.K. performed and analyzed all electrophysiology, western blot experiments, and imaging. H.K. prepared all brain samples for electron microscopy and proteomics experiments.

G.H.K. and K.J.L. performed electron microscopy and analyzed data. M.J. and S.-W.L. performed the proteomics experiments, and S.Cha and S.Chae. analyzed the proteomic data.

H.P. supervised the project. H.K., S.Chae, and H.P. wrote the main manuscript text, and all authors reviewed the manuscript.

## Methods

All mouse experiments were conducted in accordance with the ethical guidelines and protocols approved by the Institutional Animal Care and Use Committee (IACUC) of the KBRI (IACUC-25-00042).

### Mice

C57BL/6 mice (The Jackson Laboratory), Aldh1l1-CreERT2 (The Jackson Laboratory), LSL-TurboID-ER^21^ (loxP-stop-loxP-TurboID-V5-KDEL generated from Cyagen) mice, and AsTER (Aldh1l1-CreERT2;LSL-TurboID-ER) mice were used in this study. To label astrocytic proteins, Aldh1l1-CreERT2 mice were crossed with LSL-TurboID-ER mice to generate AsTER mice. All mouse lines were maintained on a C57BL/6 background. Mice were housed in standard plastic cages under specific pathogen-free conditions at the Laboratory Animal Center of the Korea Brain Research Institute. The animal facility was maintained at 20–24 °C and 40–60% relative humidity under a 12-h light/12-h dark cycle, with lights off from 20:00 to 08:00. Young adult mice aged 8–10 weeks were used for all experiments.

### Stereotaxic injection

Mice were anesthetized with isoflurane (Hana Pharm) in a sealed chamber and maintained under anesthesia using a low-flow delivery system (SomnoSuite). Stereotaxic injections were performed using a stereotaxic frame (Kopf) and a Nanojector III (Drummond Scientific) equipped with glass micropipettes (World Precision Instruments) pulled using a motorized micropipette puller (Sutter Instrument). For astrocytic CtsD knockdown and rescue, AAV-Ef1a-G-CREon-shCtsD (5.74 × 10^13^ GC/ml; generated by the Korea Institute of Science and Technology Virus Facility) and AAV-CMV-DIO-hCTSD-T2A-mScarlet (1.03 × 10^14^ GC/ml; VectorBuilder) were injected bilaterally into the dorsal hippocampus (ML: ±1.70 mm, AP: −1.9 mm from bregma, DV: −1.3 mm from the brain surface) of Aldh1l1-CreERT2 mice. A total volume of 500 nl was delivered per hemisphere in five 100-nl injections with 60-s intervals between injections. Mice were used for experiments approximately 1 month after viral injection. For pharmacological inhibition of CtsD, pepstatin A (MedChemExpress, HY-P0018) was prepared at 10 μM in PBS containing 0.1% DMSO and injected into the dorsal hippocampus at 500 nl per hemisphere. For antibody-mediated blockade of extracellular CtsD, a mouse monoclonal anti-cathepsin D antibody (Santa Cruz Biotechnology, sc-377124) or an equal mass of normal mouse IgG (Santa Cruz Biotechnology, sc-2025) was injected into the dorsal hippocampus at a dose of 1 μg per hemisphere. DiI stain (Invitrogen, D3911) was added to both preparations to visualize injection sites. To promote extracellular matrix (ECM) degradation, chondroitinase ABC (ChABC; Sigma-Aldrich, C3667) was diluted from a 50 U/ml stock solution to a final concentration of 0.1 U/ml in PBS containing 0.1% BSA. ChABC was stereotaxically injected bilaterally into the dorsal hippocampus using the coordinates described above. A total volume of 500 nl, corresponding to 0.05 mU of ChABC, was delivered per hemisphere in five 100-nl injections with 30-s intervals between injections.

### Tamoxifen and biotin injections

To induce Cre-mediated recombination, tamoxifen (Sigma-Aldrich, T5648) was dissolved in corn oil (Sigma-Aldrich, C8267) at a concentration of 10 mg/ml and stored at −20 °C. Two weeks after viral injection, mice received intraperitoneal injections of tamoxifen at 75 mg/kg body weight once daily for five consecutive days. Control mice received corn oil alone following the same injection schedule. Mice were used for subsequent experiments 2 weeks after the first injection.

To induce biotinylation of astrocytic proteins, biotin (Aladdin, B105433) was dissolved in dimethyl sulfoxide (DMSO) at a stock concentration of 48 mg/ml and diluted in PBS before injection. Starting 2 weeks after the first tamoxifen injection, AsTER mice received intraperitoneal injections of biotin at 24 mg/kg body weight or PBS once daily for five consecutive days. Subsequent experiments were performed after completion of the biotin injections.

### Cerebrospinal fluid harvest

To assess biotin labeling of astrocyte-secreted proteins in AsTER mice (Extended Data Fig. 1h, i), cerebrospinal fluid (CSF) was collected from the cisterna magna using a modified stereotaxic approach described by Shimizu et al.^60^. Mice were anesthetized by intraperitoneal injection of tribromoethanol (Avertin; Sigma-Aldrich, T48402) and fixed in a stereotaxic frame. Approximately 5–10 μl of CSF was collected per mouse using a glass capillary. CSF samples were centrifuged at 10,000 rpm for 30 s, and only samples without a visible erythrocyte pellet were used for subsequent analyses. A protease inhibitor cocktail (Quartett, PPI1015) was added to the CSF samples before western blot analysis.

### Preparation of biotinylated astrocytic proteins in hippocampal tissues

Following tamoxifen and biotin administration, AsTER mice were subjected to CFC (learning group) or maintained in their home cages (homecage control group). Mice were sacrificed 4 h after CFC or at the corresponding time point for homecage controls and transcardially perfused with PBS, after which hippocampi were isolated. Hippocampal tissues from two mice within the same experimental group were pooled for each sample to obtain sufficient material for the enrichment of biotinylated proteins. Pooled tissues were homogenized in lysis buffer (1% Triton X-100, 0.1% Tween 20, and protease inhibitor cocktail in PBS) using a bead-based homogenizer (taco™ Prep Bead Beater). Homogenates were centrifuged at 13,000 rpm for 20 min at 4 °C, and the supernatants were collected. Protein concentrations were determined using a BCA assay. Before enrichment, TurboID expression and protein biotinylation were assessed by western blotting using an anti-V5 antibody (Thermo Fisher Scientific, R960-25) and streptavidin-HRP (Thermo Fisher Scientific, 21124), respectively. Following protein quantification using a BCA assay, 5 mg of total protein from each pooled hippocampal lysate sample was used for streptavidin bead-mediated enrichment of biotinylated proteins. Each hippocampal lysate was incubated with Dynabeads M-280 Streptavidin (Invitrogen, 11205D) for 30 min at room temperature with rotation. Beads were washed five times with wash buffer (0.1% BSA, 1% Triton X-100, 0.1% Tween 20, and protease inhibitor cocktail in PBS), and bound proteins were eluted in 2% SDS at 95 °C for 3 min. Successful enrichment of biotinylated proteins was confirmed in all samples by streptavidin blotting prior to mass spectrometry analysis (Extended Data Fig. 3a, b).

### Preparation of astrocyte-conditioned medium

To identify astrocyte-secreted proteins in vitro, primary astrocyte cultures were treated with 100 μM L-glutamic acid (Sigma-Aldrich, G8415). L-Glutamic acid was prepared as a 100 mM stock solution in 1 M HCl. Conditioned media were collected from separate cultures at 15, 30, and 60 min after glutamate treatment and pooled to generate each glutamate-treated sample. Conditioned media from untreated cultures were collected as controls. Collected media were centrifuged at 3,500 rpm for 4 min at room temperature, and the supernatants were passed through a 0.22-μm syringe filter (Millex, SLGP033RB). Filtered media were concentrated using Amicon centrifugal filter units with a 10-kDa molecular weight cutoff and a 4-ml capacity (Merck, UFC801096) at 7,100 × g for 20 min at 4 °C. For buffer exchange, 50 mM Tris-HCl was added to each filter unit, followed by centrifugation under the same conditions. Concentrated samples were supplemented with a protease inhibitor cocktail and stored at −20 °C until further processing for mass spectrometry.

### Sample preparation and TMT labeling

Biotinylated proteins eluted from streptavidin beads and proteins in ACM were digested using a modified filter-aided sample preparation (FASP) method (Wiśniewski et al., 2009). Briefly, proteins were reduced in 0.1 M dithiothreitol (DTT) in 0.1 M Tris-HCl at 37 °C for 45 min with shaking and then boiled for 10 min. ACM proteins were first diluted in 8 M urea in 0.1 M Tris-HCl (pH 7.4) and reduced with 0.1 M DTT at 33 °C for 45 min, and equal amounts of protein were processed for each ACM sample. Reduced proteins were loaded onto YM-30 membrane filters (Millipore) by centrifugation at 14,000 × g and washed three times with 200 μl of 8 M urea in 0.1 M Tris-HCl (pH 8.5). Cysteines were alkylated with 50 mM iodoacetamide in 8 M urea for 25 min at room temperature in the dark, and the filters were washed four more times with 8 M urea and then with 50 mM ammonium bicarbonate (pH 8.0). Proteins were digested on the filter with sequencing-grade trypsin (Promega) at an enzyme-to-protein ratio of 1:50 (w/w) overnight at 37 °C, followed by a second digestion at 1:100 (w/w) for 6 h. Peptides were eluted by centrifugation at 14,000 × g, the filters were rinsed twice with 50 mM ammonium bicarbonate, and the combined eluates were quantified by BCA assay, vacuum-dried and stored at −80 °C. Hippocampal peptides were labeled with TMT10plex reagents (Thermo Fisher Scientific). A universal reference prepared by pooling 10 μg of peptides from every sample was labeled with channel 126. Peptides from the homecage (127N, 129N and 130N) and learning groups (127C, 130C and 129C) were labeled at 30 μg of peptides per channel, and one control sample was labeled with channel 131. TMT reagents were dissolved in anhydrous acetonitrile, added to the peptides and incubated for 1 h at room temperature, and the reaction was quenched with 5% hydroxylamine for 20 min. The labeled peptides of each set were pooled, vacuum-dried and desalted as previously described (Hyeon et al., 2023).

### LC-MS/MS analysis

TMT-labeled peptides (40 μg per set) were analyzed using a previously described dual online two-dimensional noncontiguous fractionating and concatenating reversed-phase/reversed-phase liquid chromatography (DO-2D-NCFC-RP/RPLC) system (Lee et al., 2016; Kang et al., 2022) coupled to an Orbitrap Exploris 480 mass spectrometer (Thermo Fisher Scientific). In the first dimension, peptides were separated on a mid-pH reversed-phase column (150 μm × 50 cm) with 10 mM triethylammonium bicarbonate (TEAB) in water (pH 7.5) and 10 mM TEAB in 99% acetonitrile as solvents A and B, respectively, using a 120-min gradient of 1–50% solvent B at 0.5 μl/min. The eluate was distributed into 12 fraction loops (200 μm × 22 cm) by synchronously rotating two NCFC valves every 2 min for five cycles, so that each fraction contained five noncontiguous eluates. Each fraction was acidified and diluted 10-fold with 0.2% trifluoroacetic acid, loaded onto a solid-phase extraction (SPE) column (150 μm × 3 cm), and separated on an analytical column (75 μm × 100 cm) at 300 nl/min and 60 °C with 0.1% formic acid in water and 0.1% formic acid in acetonitrile, using a gradient of 10–40% acetonitrile over 167 min, 40–80% over 5 min, and 80% for 8 min. Two SPE columns and two analytical columns were operated alternately, and all columns were packed in-house with Jupiter C18 particles (3 μm, 300 Å; Phenomenex). Peptides were ionized at 2.4 kV with a desolvation capillary temperature of 250 °C. Full MS scans (m/z 350–1,500) were acquired at a resolution of 60,000 with an automatic gain control (AGC) target of 5 × 10⁶ and a maximum injection time of 20 ms. Precursor ions were selected in a data-dependent manner with a cycle time of 1.5 s and a dynamic exclusion of 25 s, isolated with a 0.7-Th window, and fragmented by higher-energy collisional dissociation (HCD) at a normalized collision energy of 28. MS/MS spectra were acquired at a resolution of 45,000 with a fixed first mass of m/z 120, an AGC target of 1 × 10⁶, and a maximum injection time of 93 ms. ACM peptides from three untreated and three glutamate-treated cultures were analyzed individually without fractionation using a dual-online ultrahigh-pressure LC system (Lee et al., 2014; Lee et al., 2015) with two SPE and two analytical columns, as described above, coupled to the Orbitrap Exploris 480.

Peptides were separated at 300 nl/min with a 180-min gradient of 10–37.5% acetonitrile in 0.1% formic acid over 160 min, 37.5–80% over 5 min, 80% for 13 min, and 10% for 2 min. Full MS scans (m/z 350–1,400; resolution 60,000) and data-dependent MS/MS scans (resolution, 15,000; HCD normalized collision energy, 28; cycle time, 1.5 s; dynamic exclusion, 25 s; singly charged ions excluded) were acquired with AGC targets of 5 × 10⁶ and 1 × 10⁶, respectively.

### Peptide and protein identification

Precursor masses of the MS/MS spectra were refined using multiplexed post-experimental monoisotopic mass refinement (mPE-MMR; Madar et al., 2017), and the refined spectra were searched with MS-GF+ (v9949; Kim and Pevzner, 2014) against the UniProt *Mus musculus* reference proteome (63,602 entries) combined with common contaminants (179 entries).

Search parameters were a precursor mass tolerance of 10 ppm and semi-tryptic cleavage, with carbamidomethylation of cysteine (+57.0215 Da) as a fixed modification and oxidation of methionine (+15.9949 Da), deamidation of asparagine and glutamine (+0.9840 Da), and biotinylation of lysine (+226.0776 Da) as variable modifications. For the TMT data, TMT labeling (+229.1629 Da) of peptide N-termini and of lysine was additionally included, such that lysine could carry either TMT or biotin. Peptide-spectrum matches (PSMs) were filtered at a 1% false discovery rate (FDR) using the target–decoy strategy. Proteins were inferred by bipartite graph analysis (Hyeon et al., 2023): within each protein group, the protein with the largest number of sibling peptides was selected as the representative, ties were resolved by sequence coverage, and proteins with unique peptides were reported separately. Proteins identified by at least two peptides were used for further analyses.

All PSMs assigned to biotinylated peptides were manually inspected. A biotinylation site was accepted only when fragment ions flanking the modified lysine were observed and the major peaks of the spectrum were assigned. Because biotinylation of lysine prevents tryptic cleavage at its C-terminal side (Li et al., 2021), a biotin assigned to a peptide C-terminal lysine was relocated to the nearest internal lysine when the y1 ion was not supported (mass error > 10 ppm or low intensity); the PSM was discarded when no such lysine existed, unless the lysine was the C-terminus of the protein. For TMT quantification, reporter ion intensities were extracted from the MS/MS spectra and corrected for isotopic impurities.

### Differential expression and functional enrichment analysis

For the TMT data, reporter ion intensities of the PSMs were log_2_-transformed and quantile-normalized across the channels. PSMs were summed for each peptide sequence on the linear scale. To identify differentially expressed peptides between the learning and homecage groups, the log_2_-fold change of each peptide in each replicate was computed as the difference between its learning and homecage channels, and a moderated one-sample t test was applied. *P*-values were computed from an empirical null distribution obtained by enumerating all sign permutations of the pair-wise fold changes. Differentially expressed peptides were selected as the ones with *P* < 0.1 and absolute median log_2_-fold changes across replicates ≥ 0.419 (1.34-fold), corresponding to the 90^th^ percentile of the null distribution of absolute median log_2_-fold changes.

For label-free quantification of ACM proteins, an accurate mass and time (AMT) tag database was constructed from peptides identified at 1% FDR across all ACM LC-MS/MS datasets. Unique mass classes (UMCs; LC-MS features detected by mPE-MMR) that lacked an MS/MS identification in a given run were assigned peptide identities by matching to AMT tags within tolerances of ±10 ppm in mass and ±0.025 in normalized elution time (NET)^61^. Peptide abundances were calculated as the summed intensities of the MS1 components of each UMC and aligned across runs. Intensities of peptides were log_2_-transformed and quantile-normalized. Peptides detected in at least two of three replicates in both groups were tested using a moderated two-sample t test, and *P*-values were computed from an empirical null distribution obtained from all possible permutations of sample labels between the two groups. Differentially expressed peptides were selected as those with *P* < 0.1 and absolute log_2_-fold changes ≥ 0.58 (1.5-fold), together with peptides detected in at least two of three replicates in one group and in none of the other. For both datasets, differentially expressed proteins (DEPs) were defined as those with at least two differentially expressed peptides changing in the same direction and none in the opposite direction. Gene ontology (GO) enrichment analysis of the DEPs was performed using DAVID^62^. The GO terms represented by the DEPs were identified as the ones with *P* < 0.05.

### Primary astrocyte culture

Primary astrocytes were prepared using a modified AWESAM^63^ protocol. Cortical astrocytes were prepared from postnatal day 1–3 wild-type C57BL/6 mice. Pups were anesthetized by hypothermia induced by brief placement on ice, in accordance with ethical guidelines. Cortices were dissected in ice-cold Hank’s balanced salt solution (HBSS; Gibco, 14170-112), followed by incubation in pre-warmed 0.25% trypsin-EDTA at 37 °C for 20 min with gentle inversion every 5 min. After trypsinization, tissue was washed ten times with HBSS and triturated with DMEM (Hyclone, SH30243.01) supplemented with 10% heat-inactivated low-IgG fetal bovine serum (Gibco, 16250-078) and 1% penicillin-streptomycin (Gibco, 15140122) (10% DFP medium). The cell suspension was passed through a 40-μm cell strainer and plated onto 60-mm dishes (2.5 × 10⁶ cells/dish). Cultures were maintained at 37 °C in a humidified 5% CO₂ incubator. At DIV 8–10, confluent mixed glial cultures were shaken at 110 rpm for 6 h at 37 °C to remove loosely attached neurons and non-astrocytic glial cells. Cultures were then washed three times with pre-warmed PBS and detached with 0.25% trypsin-EDTA for 5 min at 37 °C, and the reaction was quenched with 10% DFP. Cells were collected by centrifugation at 3,500 rpm for 4 min at room temperature, resuspended in Neurobasal-based medium (Gibco, 21103049) containing 2% B-27 supplement (Gibco, 17504001), 2 mM GlutaMAX (Gibco, 35050061), penicillin-streptomycin, and 5 ng/ml heparin-binding epidermal growth factor (HBEGF; Sigma-Aldrich, E4643-50ug) (NB+H medium), and replated onto 0.04% polyethyleneimine (PEI; Sigma-Aldrich, P3143-100ml)-coated culture plates. PEI-coated plates were prepared by incubating plates with 0.04% PEI in 25 mM borate buffer for 1 h at 37 °C, followed by four washes with PBS. Subcultured astrocytes were maintained in NB+H medium, with half-medium changes every 2–3 days, and used within 1 week after replating.

### Lentiviral transduction and TurboID-ER labeling in primary astrocytes

To validate whether TurboID-ER can label astrocytic secretory proteins, primary astrocytes were exposed to lentiviral vectors encoding GFAP-driven TurboID-V5-KDEL-P2A-EGFP (LV-Gfap-TurboID-ER-p2A-EGFP). At DIV 8–10, lentivirus was applied to the culture medium at a multiplicity of infection (MOI) of 20 immediately before astrocyte replating. Half of the medium was replaced every 2–3 days, and transduction efficiency was monitored by EGFP fluorescence. For TurboID-ER-mediated labeling, astrocytes were incubated with 50 μM D-biotin (Aladdin, B105433) at DIV 12–14. Cells were incubated at 37 °C for 16 h and then fixed for immunocytochemistry.

### Live-cell imaging

Primary astrocytes were cultured in Confocal Dishes (SPL, 101350) and transfected with CtsD-pHluorin at DIV 12-14. Live-cell imaging was performed using a Nikon A1Rsi confocal microscope mounted on a Ti-E inverted microscope (Nikon, Japan), equipped with an environmental control system maintaining 37°C and 5% CO₂. During imaging, cells were continuously perfused with normal extracellular solution (in mM: 119 NaCl, 2.5 KCl, 20 HEPES, 2 CaCl_2_, 30 Glucose, 2 MgCl_2_). For glutamate stimulation, 100 µM glutamate was added to the normal extracellular solution. For the control group, normal extracellular solution was perfused instead of glutamate-containing solution during the stimulation period. For NH_4_Cl treatment, a modified extracellular solution was used in which 50mM NaCl was replaced with 50mM NH_4_Cl, while the concentrations of the other components remained unchanged. The pH of all solutions was adjusted to 7.4, and all solutions were prewarmed before use. Images were acquired using a 63X objective at a resolution of 512 × 512 pixels every 1 s for 7 min. Baseline fluorescence was recorded for 1 min, followed by glutamate stimulation for 3 min. Immediately after glutamate stimulation, NH_4_Cl solution was applied to alkalinize acidic compartments. Fluorescence changes were analyzed using ImageJ software and normalized as indicated in Fig. S5.

### BAPTA-AM treatment

BAPTA-AM (Sigma-Aldrich, A1076) was dissolved in DMSO and diluted in culture medium to a final concentration of 20 μM. Cells were incubated with BAPTA-AM for 30 min at 37°C, washed with fresh normal extracellular solution, and subsequently subjected to live-cell imaging. Cells treated with an equivalent volume of DMSO served as the vehicle control.

### Cell line culture and transfection

C8D1a mouse astrocytic cells were cultured in 10% DFP and incubated at 37 °C in a humidified incubator with 5% CO₂. For human cathepsin D (hCTSD) overexpression (Extended Data Fig. 14), cells were co-transfected with CMV-DIO-hCTSD-T2A-mScarlet and Ef1α-Cre plasmids using Lipofectamine 2000 (Invitrogen, 11668-019). At 48 h after transfection, cells were fixed and processed for immunocytochemistry.

### Immunocytochemistry

Primary astrocytes or C8D1a cells grown on coverslips were washed with PBS and fixed with 4% paraformaldehyde (4% PFA) for 10 min at room temperature. Following fixation, cells were washed with PBS and incubated in blocking buffer containing 5% bovine serum albumin (BSA; Sigma-Aldrich, A7906), 2% normal goat serum (Jackson Lab, 005-000-121), and 0.2% Triton X-100 (Sigma-Aldrich, X100) in PBS for 1–1.5 h at room temperature. Cells were incubated with primary antibodies diluted in blocking buffer for 1 h at room temperature. After three washes with PBS for 5 min each, cells were incubated with appropriate fluorophore-conjugated secondary antibodies diluted in blocking buffer for 1 h at room temperature. For detection of biotinylated proteins, Alexa Fluor 680-conjugated streptavidin was included during secondary antibody incubation. Coverslips were then washed with PBS and mounted on glass slides using antifade mounting medium containing DAPI (Vector Laboratories, H-1200). Details of primary and secondary antibodies, including suppliers, catalog numbers, and working dilutions, are provided in Supplementary Table 6.

### DQ-collagen IV proteolysis assay

Proteolysis of DQ-collagen IV was assessed using a modified version of the protocol described by Jedeszko et al.^44^. C8D1a cells and primary astrocytes were prepared as described above and seeded in 12-well plates coated with basement membrane extract containing DQ-collagen IV. Cultrex basement membrane extract (R&D Systems, 3432-010-01) was diluted 1:100 in ice-cold serum-free DMEM and supplemented with DQ-collagen IV (Invitrogen, D12052) at a final concentration of 25 μg/ml. Plates were coated with 600 μl of the mixture per well for 1 h at room temperature. The coating solution was then removed, and cells were seeded in culture medium.

For hCTSD overexpression, C8D1a cells were co-transfected with CMV-DIO-hCTSD-T2A-mScarlet and a Cre-expressing plasmid (hCTSD-OE group) as described above. Control C8D1a cells were transfected with EF1α-mCherry-IRES-Cre. For the hCTSD-OE + Pep group, hCTSD-overexpressing cells were treated with 10 μM pepstatin A (MedChemExpress, HY-P0018) for 1 h before fixation. Primary astrocytes were labeled with 1 μM CellTracker Orange CMTMR (Invitrogen, C2927) for 30 min at 37 °C and gently washed with culture medium. Cells were then treated with an anti-CtsD antibody (Santa Cruz Biotechnology, sc-377124; 500 ng per well) or an equal mass of normal mouse IgG (Santa Cruz Biotechnology, sc-2025), in the presence or absence of 100 μM glutamate for 1 h.

Cells were fixed with 4% paraformaldehyde for 10 min at room temperature and washed with PBS. Confocal z-stack images were acquired using a Leica STELLARIS 8 confocal microscope equipped with a 63× oil-immersion objective and LAS X software (v3.5.6.21594). Maximum-intensity projections were generated from z-stacks and analyzed using Fiji/ImageJ software, with an identical field-of-view area used across experimental groups. The same brightness and contrast settings were used for image display. Binary cell masks were generated by thresholding mCherry or mScarlet fluorescence in C8D1a cells and CellTracker Orange fluorescence in primary astrocytes using consistent thresholding criteria across groups. Individual cells were randomly selected for image analysis. Integrated DQ-collagen IV fluorescence intensity was quantified outside the cell masks, and cell area was measured from the same masks. Each z-stack contained a single cell, and no additional normalization to cell number was performed. Experiments were performed using primary cultures prepared separately from three individual mice.

### CtsD activity assay

Astrocyte-conditioned medium was prepared as described above, except that no protease inhibitor cocktail was added. CtsD activity was measured using a fluorometric CtsD activity assay kit (Abcam, ab65302) according to the manufacturer’s instructions, using equal volumes of each sample. Samples were incubated with substrate at 37 °C for 1–2 h, and fluorescence was measured using a SpectraMax iD5 microplate reader (Molecular Devices) at excitation/emission wavelengths of 328 and 460 nm, respectively. Fluorescence was recorded in relative fluorescence units (RFU). Background RFU values were subtracted from sample RFU values, and the resulting values were normalized to the mean of the untreated control group. Experiments were repeated using at least three independently prepared astrocyte cultures, each derived from a different mouse.

### Cytokine and chemokine array

To assess changes in hippocampal cytokine and chemokine levels following astrocytic CtsD knockdown, mice were transcardially perfused with PBS, and hippocampi were isolated. Isolated hippocampi were sectioned at a thickness of 400 μm in artificial cerebrospinal fluid (ACSF) using a VF-200-OZ Compresstome (Precisionary). Slices were examined under a fluorescence microscope, and only slices showing EGFP expression were selected for tissue collection. Selected tissue was homogenized in lysis buffer (1% Triton X-100, 0.1% Tween 20, protease inhibitor cocktail in PBS) using a bead-based homogenizer (taco™ Prep Bead Beater). Homogenates were centrifuged at 13,000 rpm for 20 min at 4 °C, and supernatants were collected. Protein concentrations were determined using a BCA assay, and equal amounts of protein (300 μg per membrane) were analyzed using the Proteome Profiler Mouse Cytokine Array Panel A (R&D Systems, ARY006) according to the manufacturer’s instructions. Chemiluminescent signals were acquired using a ChemiDoc MP imaging system (Bio-Rad), and cytokine and chemokine spot intensities were quantified using Fiji/ImageJ. For heatmap visualization, spot intensities were min–max normalized to a range of 0–1. Statistical comparisons between control and aCtsD-KD mice were performed using multiple unpaired t-tests, with the false discovery rate controlled at 1% using the two-stage step-up method of Benjamini, Krieger, and Yekutieli in GraphPad Prism.

### Western blotting

Protein concentrations were determined using a BCA assay kit (Thermo Scientific) according to the manufacturer’s instructions. Samples were heated in Laemmli buffer (Bio-Rad) at 95 °C for 3 min and separated on precast 4–20% polyacrylamide gels (Bio-Rad, 456-8093). Proteins were then transferred onto 0.2-μm PVDF membranes (Bio-Rad, 1704156) using a Trans-Blot Turbo Transfer system (Bio-Rad). Membranes were blocked for 1 h in 3% skim milk prepared in TBS containing 0.1% Tween 20 (TBST), followed by incubation with primary antibodies overnight at 4 °C and HRP-conjugated secondary antibodies for 2 h at room temperature. Membranes were washed with TBST after each antibody incubation. Protein bands were detected using enhanced chemiluminescence (ECL) substrates and imaged using a ChemiDoc MP Imaging System (Bio-Rad). Band intensities were quantified using Fiji/ImageJ and normalized to corresponding β-actin band intensities. Details of antibodies, working dilutions, and ECL substrates are provided in Supplementary Table 6.

### Immunohistochemistry

Mice were deeply anesthetized with 1–2% isoflurane and transcardially perfused with 1× PBS (Welgene), followed by ice-cold 4% paraformaldehyde (PFA; Biosesang) in PBS. Brains were isolated, post-fixed overnight in 4% PFA at 4 °C and cryoprotected in 30% sucrose in PBS for 48 h. Brains were then embedded in OCT compound (Leica), frozen at −80 °C, and sectioned at a thickness of 40 μm using a cryostat (Leica). For WFA staining, sections were permeabilized with 0.2% Triton X-100 in PBS and sequentially treated with streptavidin and biotin reagents using an endogenous biotin-blocking kit (Invitrogen, E21390) before the blocking step. Sections were blocked for 1 h at room temperature in PBS containing 10% goat serum and 0.2% Triton X-100, followed by incubation with primary antibodies diluted in blocking solution overnight at 4 °C. Sections were then incubated with secondary antibodies and/or fluorescently conjugated streptavidin for 2 h at room temperature. Nuclei were counterstained with DAPI (Sigma-Aldrich, D9542), and sections were mounted using antifade mounting medium (Vectashield, H-1400). Details of antibodies and other staining reagents, including working dilutions, are provided in Supplementary Table 6.

Confocal images for the analyses described below were acquired using a Leica STELLARIS 8 confocal microscope and LAS X software (v3.5.6.21594). For virus-injected mice, hippocampal tissue sections expressing EGFP (aCtsD-KD) or mScarlet (aCtsD-RE) were selected for imaging and analysis. For each staining, brightness and contrast settings were determined using negative-control sections processed for immunohistochemistry without secondary antibodies and applied identically to all corresponding images before quantification. All images were analyzed using Fiji/ImageJ software. For image-based data displayed as violin plots with dots, the violins show image-level distributions, and each dot represents the mean value for an individual mouse. These per-mouse mean values were used for statistical analysis.

To quantify cell-type-specific CtsD intensity, single optical sections were acquired using a 63× oil-immersion objective. Regions of interest (ROIs) were defined using S100β, NeuN, or Iba1 staining to identify astrocytes, neurons, or microglia, respectively. Identical brightness and contrast adjustments and thresholding settings were applied to each cell-type marker across images, and identical background subtraction settings were applied to the CtsD channel. CtsD fluorescence intensity within each ROI was measured and normalized to the corresponding marker-positive area (μm²). For each mouse, 24 images were analyzed across the CA1 pyramidal cell layer, dentate gyrus, and CA1 stratum radiatum in dorsal hippocampal tissue sections. Measurements from individual images were averaged to obtain a single value per mouse, which was then normalized to the mean value of homecage mice processed in the same staining batch.

To quantify GFAP fluorescence intensity and GFAP-positive area, z-stack images were acquired using a 63× oil-immersion objective lens. Each z-stack consisted of 10 optical sections acquired at 1-μm intervals, and maximum-intensity projections were generated for quantification. Identical thresholding settings were applied to the GFAP channel. Mean GFAP fluorescence intensity was measured across the entire field of view, and GFAP-positive area was quantified within threshold-defined regions. For each mouse, 16 image stacks from dorsal hippocampal tissue sections were analyzed. Measurements from individual images were averaged to obtain a single value per mouse.

To quantify WFA fluorescence intensity and WFA-positive area, z-stack images were acquired using a 63× oil-immersion objective lens. Each z-stack consisted of 10 optical sections acquired at 1-μm intervals, and maximum-intensity projections were generated for quantification. Identical thresholding settings were applied to the WFA channel. Mean WFA fluorescence intensity was measured across the entire field of view, and WFA-positive area was quantified within threshold-defined regions. For each mouse, 8 image stacks from dorsal hippocampal tissue sections were analyzed, and measurements were averaged for each mouse. For ChABC dose optimization, WFA fluorescence intensity in the ChABC-injected hemisphere was normalized to that in the contralateral PBS-injected hemisphere and expressed as the ChABC/PBS intensity ratio.

To quantify glial lysosomal density and ECM engulfment, z-stack images were acquired using a 63× oil-immersion objective lens. Each z-stack consisted of 10 optical sections acquired at 1-μm intervals, and a single optical section containing clearly identifiable glial cell bodies and processes was selected from each stack for quantification. The DiAna plugin was used to count LAMP1+ puncta colocalized with S100β or Iba1 and LAMP1+ puncta colocalized with both WFA and the corresponding glial marker. Puncta counts were normalized to the corresponding marker-positive area and expressed per 10 μm². S100β+ and Iba1+ areas were measured in the same optical sections using identical thresholding settings for each marker. For each mouse, 8 images from dorsal hippocampal tissue sections were analyzed, and measurements were averaged for each mouse.

### Whole-cell patch clamp

Following isoflurane anesthesia, mice were transcardially perfused with 15–20 ml of ice-cold, oxygenated slicing ACSF (in mM: 93 *N*-methyl-D-glutamine [NMDG], 93 HCl, 2.5 KCl, 1.2 NaH₂PO₄, 30 NaHCO₃, 20 HEPES, 5 sodium ascorbate, 2 thiourea, 3 sodium pyruvate, 12 *N*-acetyl-L-cysteine [NAC], 0.5 CaCl₂, 10 MgCl₂, and 25 glucose; pH 7.3–7.4). Brains were removed and submerged in the same solution, and 400-μm-thick horizontal slices containing the hippocampus were prepared using a Compresstome (VF-200-OZ, Precisionary). Slices were allowed to recover at 30–32 °C for 50 min in recovery ACSF (in mM: 104 NaCl, 2.5 KCl, 1.2 NaH₂PO₄, 24 NaHCO₃, 5 HEPES, 5 sodium ascorbate, 2 thiourea, 3 sodium pyruvate, 12 NAC, 2 CaCl₂, 2 MgCl₂, and 13 glucose; pH 7.3–7.4) before recording. After recovery, slices were transferred to a recording chamber and continuously perfused with oxygenated recording ACSF (in mM: 124 NaCl, 2.5 KCl, 1.2 NaH₂PO₄, 24 NaHCO₃, 5 HEPES, 2 CaCl₂, 2 MgCl₂, and 13 glucose; pH 7.3–7.4) at 2–3 ml/min. Recordings were performed at 20–25 °C.

Whole-cell recordings were obtained from CA1 pyramidal neurons using borosilicate glass micropipettes (3–5 MΩ). Two internal solutions were used: a cesium methanesulfonate (CsMeSO₃)-based solution (in mM: 140 CsMeSO₃, 7 NaCl, 0.2 EGTA, 2 MgCl₂, 4 Mg-ATP, 0.3 Na₂-GTP, 10 Na₂-phosphocreatine, and 10 HEPES) and a CsCl-based solution (in mM: 140 CsCl, 7 NaCl, 0.2 EGTA, 2 MgCl₂, 4 Mg-ATP, 0.3 Na₂-GTP, 10 Na₂-phosphocreatine, 10 HEPES, and 1 QX-314). Both internal solutions had a pH of 7.3 and an osmolality of 290–300 mOsm.

For recordings in astrocytic CtsD-knockdown mice, CA1 pyramidal neurons were selected within the virally transduced region near EGFP-expressing astrocytes. sEPSCs were recorded at −70 mV using pipettes filled with CsMeSO₃-based solution, with 10 μM bicuculline (Tocris) added to the recording ACSF to block GABAergic transmission. sIPSCs were recorded at −70 mV using pipettes filled with CsCl-based solution, with 10 μM NBQX (Tocris) and 50 μM D-AP5 (Tocris) added to the recording ACSF to block glutamatergic transmission. To evoke synaptic responses, a bipolar stimulation electrode (CBBPE75, FHC) was placed in CA1 stratum radiatum to stimulate Schaffer collaterals. eEPSCs, eIPSCs, and paired-pulse responses were recorded from the same CA1 pyramidal neuron using a pipette filled with CsMeSO₃-based solution. eEPSCs and paired-pulse responses were recorded at a holding potential of −70 mV, whereas eIPSCs were recorded at 0 mV. The EPSC/IPSC ratio was calculated as the eEPSC amplitude divided by the eIPSC amplitude recorded from the same neuron. Paired-pulse ratios were measured at interstimulus intervals of 20, 50, and 100 ms and calculated as the amplitude of the second EPSC divided by that of the first. eEPSCs and eIPSCs were validated in independent experiments by application of NBQX plus D-AP5 and bicuculline, respectively (data not shown).

For acute antibody-mediated blockade of extracellular CtsD, slices from wild-type mice were preincubated in recording ACSF containing 500 ng/ml anti-CtsD antibody (Santa Cruz Biotechnology, sc-377124) or 500 ng/ml normal mouse IgG (Santa Cruz Biotechnology, sc-2025) for 30 min before recording. The respective antibodies were maintained at the same concentration throughout recording. sEPSCs and sIPSCs were recorded sequentially from the same CA1 pyramidal neuron at −70 and 0 mV, respectively, using CsMeSO₃-based internal solution.

Data were acquired using Clampex 10.6 software (Axon Instruments) and a MultiClamp 700B amplifier (Molecular Devices), low-pass filtered at 10 kHz, and digitized at 10–50 kHz. All whole-cell patch-clamp data were analyzed using Clampfit 11.2 (Axon Instruments).

### Extracellular fEPSP recording

Following isoflurane anesthesia, brains were removed from mice and submerged in ice-cold, oxygenated sucrose-based slicing ACSF (in mM: 5 KCl, 1.25 NaH₂PO₄, 26 NaHCO₃, 0.5 CaCl₂, 10 MgCl₂, 10 D-glucose, and 212.5 sucrose; pH 7.3–7.4). Hippocampi were isolated and cut into 400-μm-thick transverse slices in the same solution using a Compresstome (VF-200-OZ, Precisionary). Slices were allowed to recover at 27 °C for 50 min in standard ACSF (in mM: 124 NaCl, 5 KCl, 1.25 NaH₂PO₄, 26 NaHCO₃, 10 D-glucose, 2.5 CaCl₂, and 1.5 MgCl₂; pH 7.3–7.4) before recording. After recovery, slices were transferred to a recording chamber and continuously perfused with oxygenated standard ACSF at 2–3 ml/min. For pharmacological experiments, slices were preincubated in standard ACSF containing 100 μM pepstatin A (MedChemExpress, HY-P0018), 500 ng/ml anti-CtsD antibody (Santa Cruz Biotechnology, sc-377124), or 500 ng/ml normal mouse IgG (Santa Cruz Biotechnology, sc-2025) for 30 min before recording. Respective agents were maintained at the same concentrations throughout all recordings, including input–output measurements, baseline recording, and LTP induction and monitoring.

For CA3–CA1 Schaffer collateral pathway recording, fEPSPs were recorded from CA1 stratum radiatum using glass micropipettes (2–3 MΩ) filled with standard ACSF, with a stimulation electrode placed on Schaffer collaterals. Stimulus intensity was set to 40–50% of maximal fEPSP slope, as determined from input–output curves plotting fEPSP slope versus stimulus intensity. After recording at least 20 min of stable baseline fEPSP responses, LTP was induced by high-frequency stimulation (HFS; 100 Hz, 1 train of 100 stimuli for 1 s) delivered through a tungsten bipolar electrode placed on Schaffer collaterals.

The fEPSP slopes were analyzed using WinLTP software. PTP was determined by averaging fEPSP slopes during the first 5 min immediately after HFS and normalizing to the baseline average. LTP was quantified by calculating the mean fEPSP slope 160–180 min post-HFS, expressed as a percentage of the 20-min baseline period.

### Transmission electron microscopy (TEM)

Control (n = 4) and aCtsD-KD (n = 4) mice were deeply anesthetized with isoflurane and transcardially perfused with 0.15 M cacodylate buffer (pH 7.4), followed by 2% paraformaldehyde and 2.5% glutaraldehyde in 0.15 M cacodylate buffer. Brains were removed, post-fixed overnight at 4 °C in the same fixative, and sectioned coronally at a thickness of 150 μm using a vibratome (VT1000S, Leica Microsystems). Tissue blocks containing the hippocampal CA1 region with fluorescently labeled astrocytes were microdissected from each slice and processed for TEM. To improve the contrast of PSDs for quantitative ultrastructural analysis, tissue blocks were washed in 0.15 M cacodylate buffer, post-fixed in 2% osmium tetroxide (18459, Ted Pella) for 1 h, and stained en bloc with 1% uranyl acetate overnight at 4 °C. Samples were then dehydrated through a graded ethanol series, infiltrated with acetone, embedded in Epon 812 resin (41420, EMS), and polymerized at 60 °C for 48 h. Ultrathin sections (80 nm) were cut using an ultramicrotome (EM UC7, Leica Microsystems), collected on 200-mesh nickel grids, and post-stained with Uranyless (22409, EMS) and lead citrate (22410, EMS). Images were acquired at 3,500× magnification using a Tecnai 20 transmission electron microscope (Thermo Fisher Scientific).

Excitatory synapses, inhibitory synapses, and MSBs were quantified using a systematic random sampling strategy, with an unbiased counting frame (5 μm × 5 μm) placed within the CA1 region. Synapses were identified according to established ultrastructural criteria: excitatory synapses were characterized by asymmetric PSDs with round presynaptic vesicles, and inhibitory synapses were characterized by symmetric PSD specializations with flattened or pleomorphic vesicles. MSBs were defined as single presynaptic boutons contacting two or more asymmetric PSDs. All images were analyzed using Fiji/ImageJ software.

### Automated tape-collecting ultramicrotome scanning electron microscopy (ATUM-SEM)

To reconstruct dendritic spines and presynaptic boutons surrounding fluorescently labeled astrocytes in three dimensions, brains from control (n = 3) and aCtsD-KD (n = 3) mice prepared as described above were processed for ATUM-SEM. Following fixation, 150-μm-thick coronal sections containing the hippocampal CA1 region were imaged by confocal microscopy (A1R, Nikon), and tissue blocks containing fluorescently labeled astrocytes were microdissected.

Tissue blocks were rinsed in 0.15 M cacodylate buffer and processed using a sequential heavy metal staining protocol. Briefly, samples were treated with 2% osmium tetroxide/1.5% potassium ferrocyanide for 1 h, incubated in 1% thiocarbohydrazide for 20 min, and subjected to a second 2% osmium tetroxide treatment for 30 min, with distilled water washes between each step. Samples were then stained en bloc with 1% aqueous uranyl acetate overnight at 4 °C and lead aspartate for 30 min at 60 °C. After dehydration through a graded ethanol series and acetone infiltration, tissues were embedded in Epon 812 resin containing 7% conductive carbon powder on aluminum rivets and polymerized at 60 °C for 48 h.

Serial ultrathin sections (50 nm) were collected onto carbon nanotube-coated polyethylene terephthalate tape (Boeckeler Instruments) using an ATUMtome (Boeckeler Instruments) and mounted onto aluminum wafers. Sections were imaged using a Sigma SEM (Carl Zeiss) equipped with a backscattered electron detector. High-resolution serial images were acquired at a pixel size of 5 × 5 nm (5 kV, 7 μs dwell time) from approximately 150 consecutive sections, corresponding to a reconstructed depth of ∼7.5 μm.

For quantitative ultrastructural analysis, CA1 pyramidal neuron dendrites, their associated dendritic spines and PSDs, and nearby axons were identified within regions located approximately 5–10 μm from EGFP-labeled astrocyte somata in each serial image stack. These structures were reconstructed in three dimensions using RECONSTRUCT software (https://synapseweb.clm.utexas.edu/software-0). Additionally, mitochondria and synaptic vesicles within presynaptic boutons were segmented to enable quantitative analysis of their ultrastructural features.

### Open field test

The open field test was performed to assess locomotor activity and exploration of central and peripheral zones. Mice were individually placed in the center of a square arena (30 × 30 × 25 cm; length × width × height) and allowed to explore freely for 5 min. Behavior was recorded using EthoVision 3.0 (Noldus), and video files were analyzed using SMART 3.0 (Panlab) to quantify total distance traveled, mean movement speed, and time spent in central and peripheral zones. After each trial, the arena was cleaned with 70% ethanol.

### Contextual fear conditioning test

The CFC test was performed in sound-attenuating chambers equipped with electrified grid floors (Panlab). During training, mice were allowed to explore the chamber for 3 min before receiving three foot shocks (0.5 mA, 2 s each), separated by 30-s intervals. Mice remained in the chamber for 1 min after the final shock before being returned to their home cages. Contextual memory was assessed 24 h after training by returning mice to the same chamber for 3 min without foot shocks. Behavior was recorded using a camera, and freezing was automatically quantified using a load-cell-based detection system and Packwin software (Panlab).

### Novel object location test

The novel object location test was performed in the arena used for the open field test, with a striped visual cue attached to one wall. Mice were habituated to the empty arena for 5 min on the first day. On the following day, mice were allowed to explore two identical objects for 5 min. After 24 h, one object was moved to a new location, and mice were returned to the arena for a 5-min memory test. The discrimination index (DI) was calculated as the ratio of the difference in exploration time between the relocated (novel) and stationary (familiar) objects to the total exploration time. Mice with a total object exploration time of less than 5 s during the learning session were excluded from analysis.

*DI* = (*T*_novel_ − *T*_familiar_) / (*T*_novel_ + *T*_familiar_)

### Statistics

All statistical analyses were performed using GraphPad Prism 11 with 95% confidence intervals, and values are expressed as the mean ± SEM. Comparisons between two groups were performed using an unpaired two-tailed Student’s *t*-test, whereas comparisons among three or more groups were performed using one-way ANOVA followed by Tukey’s multiple-comparisons test. Cumulative distributions of dendritic spine volume and PSD length were compared using the two-sample Kolmogorov–Smirnov test. Statistical analyses of data presented as violin plots and individual dots were performed on a per-mouse basis. Relationships between freezing levels and normalized CtsD intensity in astrocytes, neurons, and microglia were assessed using simple linear regression. Input–output relationships, paired-pulse ratios, and freezing levels during fear conditioning training were analyzed using two-way repeated-measures ANOVA followed by Tukey’s multiple-comparisons test. Statistical tests for each experiment are described in the respective figure legends. *P*-values less than 0.05 were considered statistically significant. n.s. indicates not significant. All experiments were independently repeated at least three times.

## Data availability statement

The mass spectrometry proteomics data have been deposited to the ProteomeXchange Consortium via the PRIDE partner repository with the dataset identifier PXD086233, and the data were also deposited in K-BDS (Korea BioData Station, https://kbds.re.kr) with the accession ID KAP242842.

## Declaration of generative AI and AI-assisted technologies in the manuscript preparation process

During the preparation of this work, the author(s) used Claude and Gemini for for proofreading and grammar checking of the manuscript text but not for data analysis, figure generation, or the generation or interpretation of scientific content. The author(s) reviewed all LLM-suggested edits and take full responsibility for the content of the published article.

## Notes

### Competing Interest Statement

The authors have declared no competing interest.

## References

1. Allen, N. J. & Eroglu, C. Cell Biology of Astrocyte-Synapse Interactions. Neuron 96, 697–708 (2017).

2. Ullian, E. M., Sapperstein, S. K., Christopherson, K. S. & Barres, B. A. Control of synapse number by glia. Science 291, 657–61 (2001).

3. Raghunathan, K. & Eroglu, C. Developmental roles of astrocytes in circuit wiring. Curr. Opin. Neurobiol. 92, 103042 (2025).

4. Christopherson, K. S. et al. Thrombospondins are astrocyte-secreted proteins that promote CNS synaptogenesis. Cell 120, 421–433 (2005).

5. Kucukdereli, H. et al. Control of excitatory CNS synaptogenesis by astrocyte-secreted proteins Hevin and SPARC. Proc. Natl. Acad. Sci. U. S. A. 108, E440–9 (2011).

6. Allen, N. J. et al. Astrocyte glypicans 4 and 6 promote formation of excitatory synapses via GluA1 AMPA receptors. Nature 486, 410–414 (2012).

7. Wang, Y. et al. Astrocyte-secreted IL-33 mediates homeostatic synaptic plasticity in the adult hippocampus. Proc. Natl. Acad. Sci. U. S. A. 118, (2021).

8. Elmariah, S. B., Oh, E. J., Hughes, E. G. & Balice-Gordon, R. J. Astrocytes regulate inhibitory synapse formation via Trk-mediated modulation of postsynaptic GABAA receptors. J. Neurosci. 25, 3638–50 (2005).

9. Irala, D. et al. Astrocyte-secreted neurocan controls inhibitory synapse formation and function. Neuron 112, 1657–1675.e10 (2024).

10. Dityatev, A. & Schachner, M. Extracellular matrix molecules and synaptic plasticity. Nat. Rev. Neurosci. 4, 456–468 (2003).

11. Fawcett, J. W., Oohashi, T. & Pizzorusso, T. The roles of perineuronal nets and the perinodal extracellular matrix in neuronal function. Nat. Rev. Neurosci. 20, 451–465 (2019).

12. Zhu, H., Li, Q., Li, Y. & Wang, W. Extracellular matrix and perineuronal nets in neural plasticity, memory, and epileptogenesis. Front. Mol. Neurosci. 19, 1945520 (2026).

13. Wang, X. et al. Extracellular proteolysis by matrix metalloproteinase-9 drives dendritic spine enlargement and long-term potentiation coordinately. Proc. Natl. Acad. Sci. U. S. A. 105, 19520–5 (2008).

14. Singh, J. B. et al. Activity-dependent extracellular proteolytic cascade cleaves the ECM component brevican to promote structural plasticity. EMBO Rep. 27, 163–185 (2026).

15. Nguyen, P. T. et al. Microglial Remodeling of the Extracellular Matrix Promotes Synapse Plasticity. Cell 182, 388–403.e15 (2020).

16. Nakajo, H. et al. Extracellular matrix proteolysis maintains synapse plasticity during brain development. Nat. Neurosci. 29, 567–580 (2026).

17. Wiese, S., Karus, M. & Faissner, A. Astrocytes as a source for extracellular matrix molecules and cytokines. Front. Pharmacol. 3, 120 (2012).

18. Branon, T. C. et al. Efficient proximity labeling in living cells and organisms with TurboID. Nat. Biotechnol. 36, 880–887 (2018).

19. Benes, P., Vetvicka, V. & Fusek, M. Cathepsin D--many functions of one aspartic protease. Crit. Rev. Oncol. Hematol. 68, 12–28 (2008).

20. Kim, H. et al. Serpina1e mediates the exercise-induced enhancement of hippocampal memory in male mice. Nat. Commun. 17, 5018 (2026).

21. Hernandez, P. J. & Abel, T. The role of protein synthesis in memory consolidation: Progress amid decades of debate. Neurobiol. Learn. Mem. 89, 293–311 (2008).

22. Freeman, F. M., Rose, S. P. R. & Scholey, A. B. Two time windows of anisomycin-induced amnesia for passive avoidance training in the day-old chick. Neurobiol. Learn. Mem. 63, 291–295 (1995).

23. Quevedo, J. et al. Two time windows of anisomycin-induced amnesia for inhibitory avoidance training in rats: protection from amnesia by pretraining but not pre-exposure to the task apparatus. Learn. Mem. 6, 600–607 (1999).

24. Greco, T. M., Seeholzer, S. H., Mak, A., Spruce, L. & Ischiropoulos, H. Quantitative mass spectrometry-based proteomics reveals the dynamic range of primary mouse astrocyte protein secretion. J. Proteome Res. 9, 2764–74 (2010).

25. Uhlén, M. et al. The human secretome. Sci. Signal. 12, (2019).

26. Mahley, R. W. Apolipoprotein E: cholesterol transport protein with expanding role in cell biology. Science 240, 622–30 (1988).

27. Humphreys, D. T., Carver, J. A., Easterbrook-Smith, S. B. & Wilson, M. R. Clusterin has chaperone-like activity similar to that of small heat shock proteins. J. Biol. Chem. 274, 6875–81 (1999).

28. Corder, E. H. et al. Gene dose of apolipoprotein E type 4 allele and the risk of Alzheimer’s disease in late onset families. Science 261, 921–3 (1993).

29. Harold, D. et al. Genome-wide association study identifies variants at CLU and PICALM associated with Alzheimer’s disease. Nat. Genet. 41, 1088–93 (2009).

30. Lin, P. et al. Calnuc binds to Alzheimer’s beta-amyloid precursor protein and affects its biogenesis. J. Neurochem. 100, 1505–1514 (2007)

31. Bonito-Oliva, A., Barbash, S., Sakmar, T. P. & Graham, W. V. Nucleobindin 1 binds to multiple types of pre-fibrillar amyloid and inhibits fibrillization. Sci. Rep. 7, 42880 (2017).

32. Stoka, V., Turk, V. & Turk, B. Lysosomal cathepsins and their regulation in aging and neurodegeneration. Ageing Res. Rev. 32, 22–37 (2016).

33. Siintola, E. et al. Cathepsin D deficiency underlies congenital human neuronal ceroid-lipofuscinosis. Brain 129, 1438–1445 (2006).

34. Koike, M. et al. Cathepsin D deficiency induces lysosomal storage with ceroid lipofuscin in mouse CNS neurons. J. Neurosci. 20, 6898–906 (2000).

35. Matsuda, N. et al. Differential activity-dependent secretion of brain-derived neurotrophic factor from axon and dendrite. J. Neurosci. 29, 14185–14198 (2009).

36. Park, H., Popescu, A. & Poo, M.-M. Essential role of presynaptic NMDA receptors in activity-dependent BDNF secretion and corticostriatal LTP. Neuron 84, 1009–1022 (2014).

37. Kim, S.-C., Kim, A., Park, J.-Y. & Hwang, E. M. Improved AAV vector system for cell-type-specific RNA interference. J. Neurosci. Methods 368, 109452 (2022).

38. Dityatev, A., Schachner, M. & Sonderegger, P. The dual role of the extracellular matrix in synaptic plasticity and homeostasis. Nat. Rev. Neurosci. 11, 735–46 (2010).

39. Ruiz-Blázquez, P. et al. Cathepsin D is essential for the degradomic shift of macrophages required to resolve liver fibrosis. Mol. Metab. 87, 101989 (2024).

40. Alcaraz, L. B. et al. A 9-kDa matricellular SPARC fragment released by cathepsin D exhibits pro-tumor activity in the triple-negative breast cancer microenvironment. Theranostics 11, 6173–6192 (2021).

41. Jedeszko, C., Sameni, M., Olive, M. B., Moin, K. & Sloane, B. F. Visualizing protease activity in living cells: from two dimensions to four dimensions. Curr. Protoc. Cell Biol. Chapter 4, Unit 4.20 (2008).

42. Briozzo, P., Morisset, M., Capony, F., Rougeot, C. & Rochefort, H. In vitro degradation of extracellular matrix with Mr 52,000 cathepsin D secreted by breast cancer cells. Cancer Res. 48, 3688–92 (1988).

43. Balian, G., Click, E. M., Crouch, E., Davidson, J. M. & Bornstein, P. Isolation of a collagen-binding fragment from fibronectin and cold-insoluble globulin. Journal of Biological Chemistry 254, 1429–1432 (1979).

44. Handley, C. J. et al. Cathepsin D cleaves aggrecan at unique sites within the interglobular domain and chondroitin sulfate attachment regions that are also cleaved when cartilage is maintained at acid pH. Matrix Biol. 20, 543–53 (2001).

45. van der Stappen, J. W., Williams, A. C., Maciewicz, R. A. & Paraskeva, C. Activation of cathepsin B, secreted by a colorectal cancer cell line requires low pH and is mediated by cathepsin D. Int. J. Cancer 67, 547–54 (1996).

46. Laurent-Matha, V. et al. Proteolysis of cystatin C by cathepsin D in the breast cancer microenvironment. FASEB J. 26, 5172–81 (2012).

47. Maynadier, M. et al. Cathepsin D stimulates the activities of secreted plasminogen activators in the breast cancer acidic environment. Int. J. Oncol. 43, 1683–90 (2013).

48. Sapolsky, A. I., Howell, D. S. & Woessner, J. F. Neutral proteases and cathepsin D in human articular cartilage. J. Clin. Invest. 53, 1044–53 (1974).

49. Chesler, M. Regulation and modulation of pH in the brain. Physiol. Rev. 83, 1183–221 (2003).

50. Du, J. et al. Protons are a neurotransmitter that regulates synaptic plasticity in the lateral amygdala. Proc. Natl. Acad. Sci. U. S. A. 111, 8961–6 (2014).

51. Fawcett, J. W. et al. The extracellular matrix and perineuronal nets in memory. Mol. Psychiatry 27, 3192–3203 (2022).

52. de Vivo, L. et al. Extracellular matrix inhibits structural and functional plasticity of dendritic spines in the adult visual cortex. Nat. Commun. 4, 1484 (2013).

53. Orlando, C., Ster, J., Gerber, U., Fawcett, J. W. & Raineteau, O. Perisynaptic Chondroitin Sulfate Proteoglycans Restrict Structural Plasticity in an Integrin-Dependent Manner. The Journal of Neuroscience 32, 18009–18017 (2012).

54. Chavis, P. & Westbrook, G. Integrins mediate functional pre- and postsynaptic maturation at a hippocampal synapse. Nature 411, 317–21 (2001).

55. Vainchtein, I. D. et al. Astrocyte-derived interleukin-33 promotes microglial synapse engulfment and neural circuit development. Science 359, 1269–1273 (2018).

56. Yu, S.-S. et al. M1-type microglia can induce astrocytes to deposit chondroitin sulfate proteoglycan after spinal cord injury. Neural Regen. Res. 17, 1072–1079 (2022).

57. Vidoni, C., Follo, C., Savino, M., Melone, M. A. B. & Isidoro, C. The Role of Cathepsin D in the Pathogenesis of Human Neurodegenerative Disorders. Med. Res. Rev. 36, 845–70 (2016).

58. Végh, M. J. et al. Reducing hippocampal extracellular matrix reverses early memory deficits in a mouse model of Alzheimer’s disease. Acta Neuropathol. Commun. 2, 76 (2014).

59. Végh, M. J. et al. Hippocampal extracellular matrix levels and stochasticity in synaptic protein expression increase with age and are associated with age-dependent cognitive decline. Mol. Cell. Proteomics 13, 2975–85 (2014).

60. Shimizu, K., Gupta, A., Brastianos, P. K. & Wakimoto, H. Anatomy-oriented stereotactic approach to cerebrospinal fluid collection in mice. Brain Res. 1774, 147706 (2022).

61. Jaitly, N. et al. Robust algorithm for alignment of liquid chromatography–mass spectrometry analyses in an accurate mass and time tag data analysis pipeline. Anal. Chem. 78, 7397–7409 (2006).

62. Sherman, B. T. et al. DAVID: a web server for functional enrichment analysis and functional annotation of gene lists (2021 update). Nucleic Acids Res. 50, W216–W221 (2022).

63. Wolfes, A. C. et al. A novel method for culturing stellate astrocytes reveals spatially distinct Ca2+ signaling and vesicle recycling in astrocytic processes. J. Gen. Physiol. 149, 149–170 (2017).

