## Supplemental Data 1 for "Astrocyte-secreted Cathepsin D regulates hippocampal memory through extracellular matrix remodeling"

This file contains supplementary Fig S1–15 and table S5, S6.

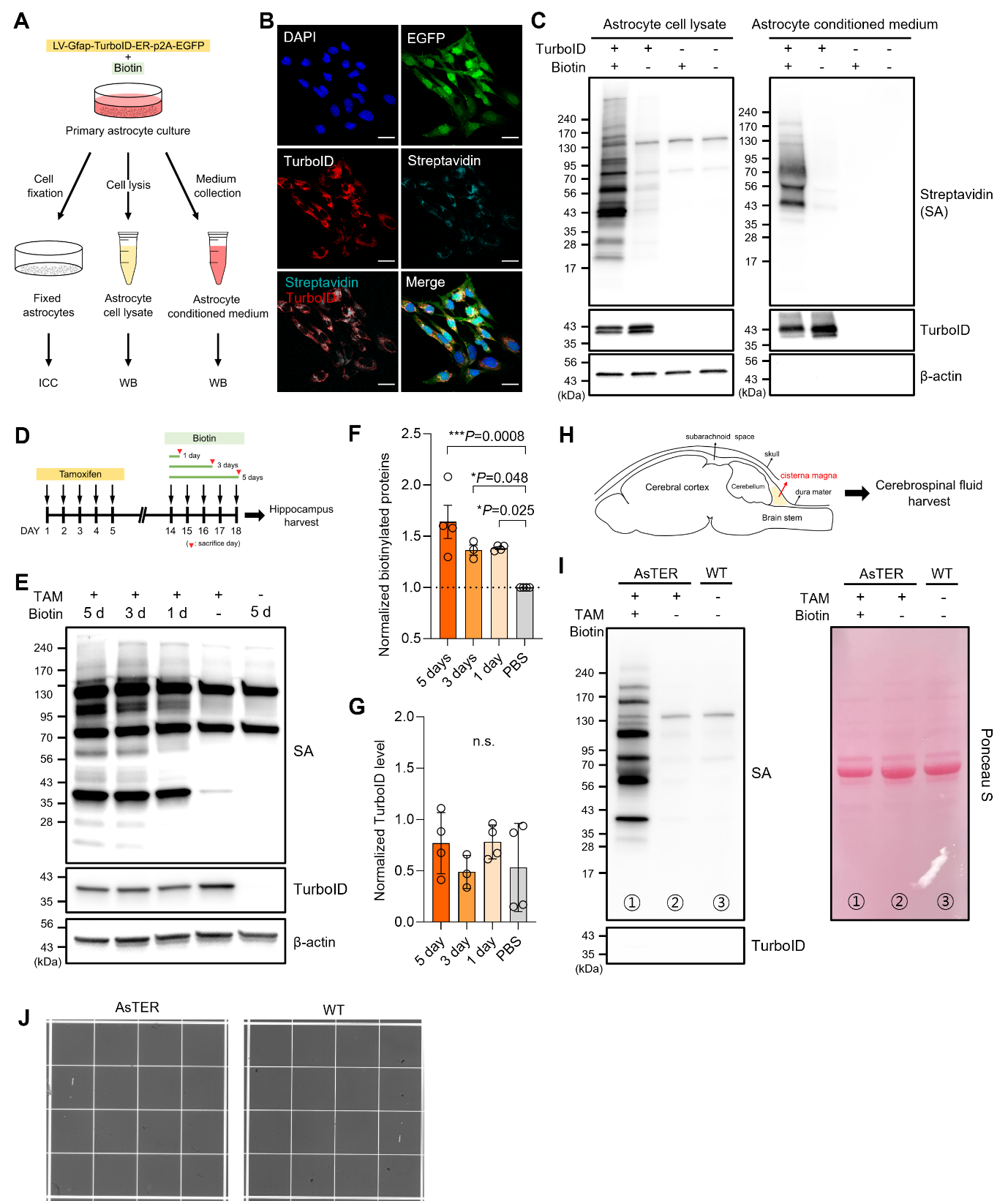

**Figure S1. TurboID-ER enables selective labeling of astrocytic proteins *in vitro* and *in vivo.***

**A.** Schematic illustration of the primary astrocyte cultures transduced with LV-Gfap-TurboID-ER-p2A-EGFP and subsequently treated with biotin. Astrocyte lysates and conditioned media were collected for western blotting (WB), or cells were fixed for immunocytochemistry (ICC). **B.** Representative confocal images of primary cultured astrocytes transduced with LV-Gfap-TurboID-ER-p2A-EGFP and labeled with streptavidin. Scale bar, 20 μm. **C.** Representative western blot images of astrocyte cell lysates and conditioned media from the cultures shown in (a). Biotinylated proteins were detected using streptavidin. **D.** Experimental timeline for *in vivo* biotinylation in AsTER mice. Biotin was administered daily for 1, 3, or 5 days, and hippocampi were collected on the indicated day (red arrowhead). **E.** Representative western blot images of hippocampal lysates from AsTER mice following biotin administration for the indicated durations. **F.** Quantification of biotinylated protein levels normalized to β-actin. **G.** Quantification of TurboID levels normalized to β-actin. **F, G.** One-way ANOVA followed by Dunnett’s multiple-comparisons test. 5 days, n = 4 mice; 3 days, n = 3 mice; 1 day, n = 4 mice; PBS, n = 4 mice. **H.** Schematic illustration of cerebrospinal fluid (CSF) collection. **I.** Representative western blot and Ponceau S staining images showing biotinylated astrocyte-derived proteins in CSF collected from AsTER and wild-type mice. **J.** Representative images of red blood cell counting to assess blood contamination in the CSF samples, showing no detectable red blood cells.

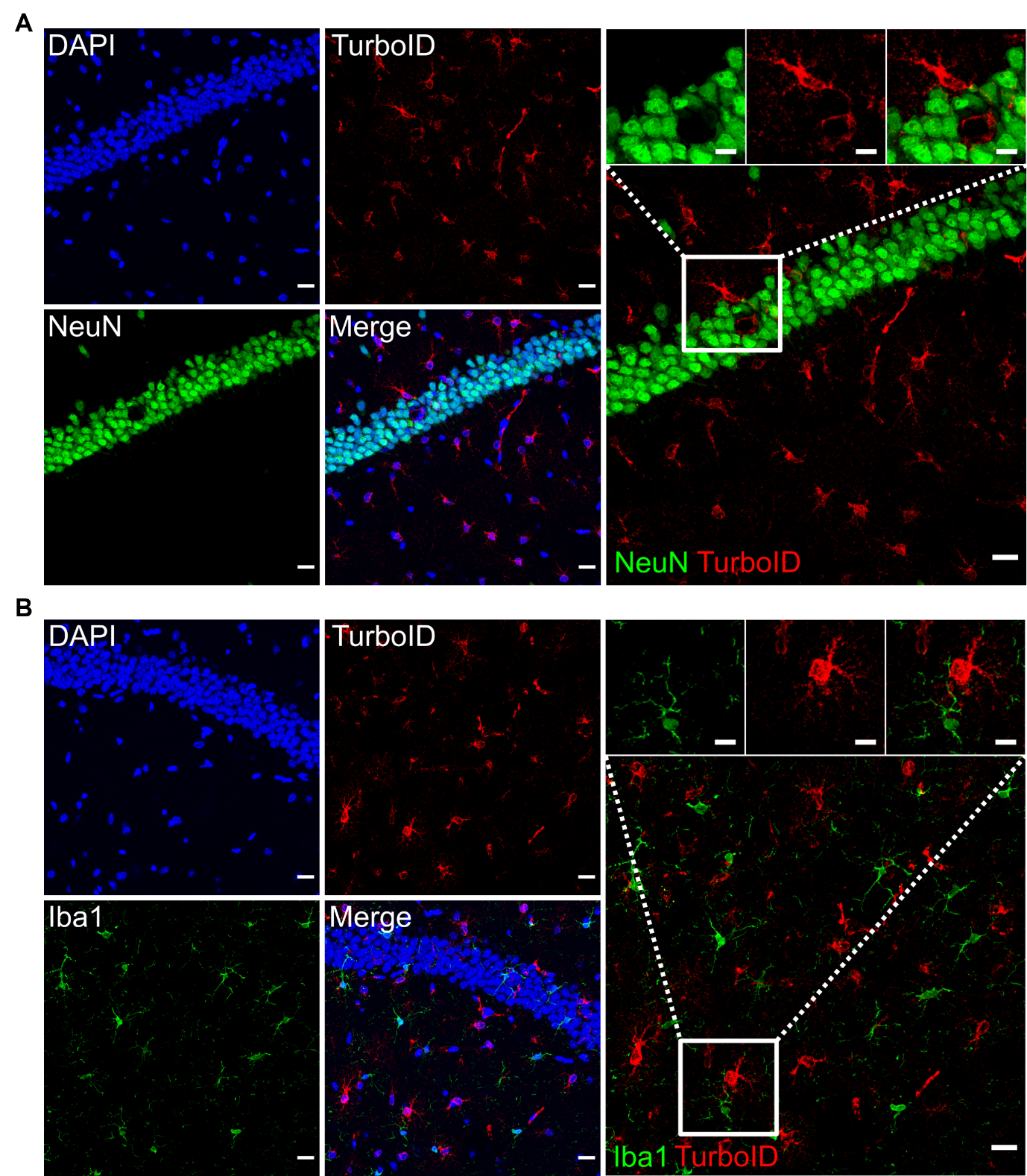
**Figure S2. Validation of astrocyte-specific TurboID expression.**

**A, B**. Representative confocal images of the hippocampal CA1 region stained for TurboID-ER (TurboID) and neuronal (NeuN; **A**) or microglial (Iba1; **B**) markers. Scale bar, 20 μm; insets, 5 μm.

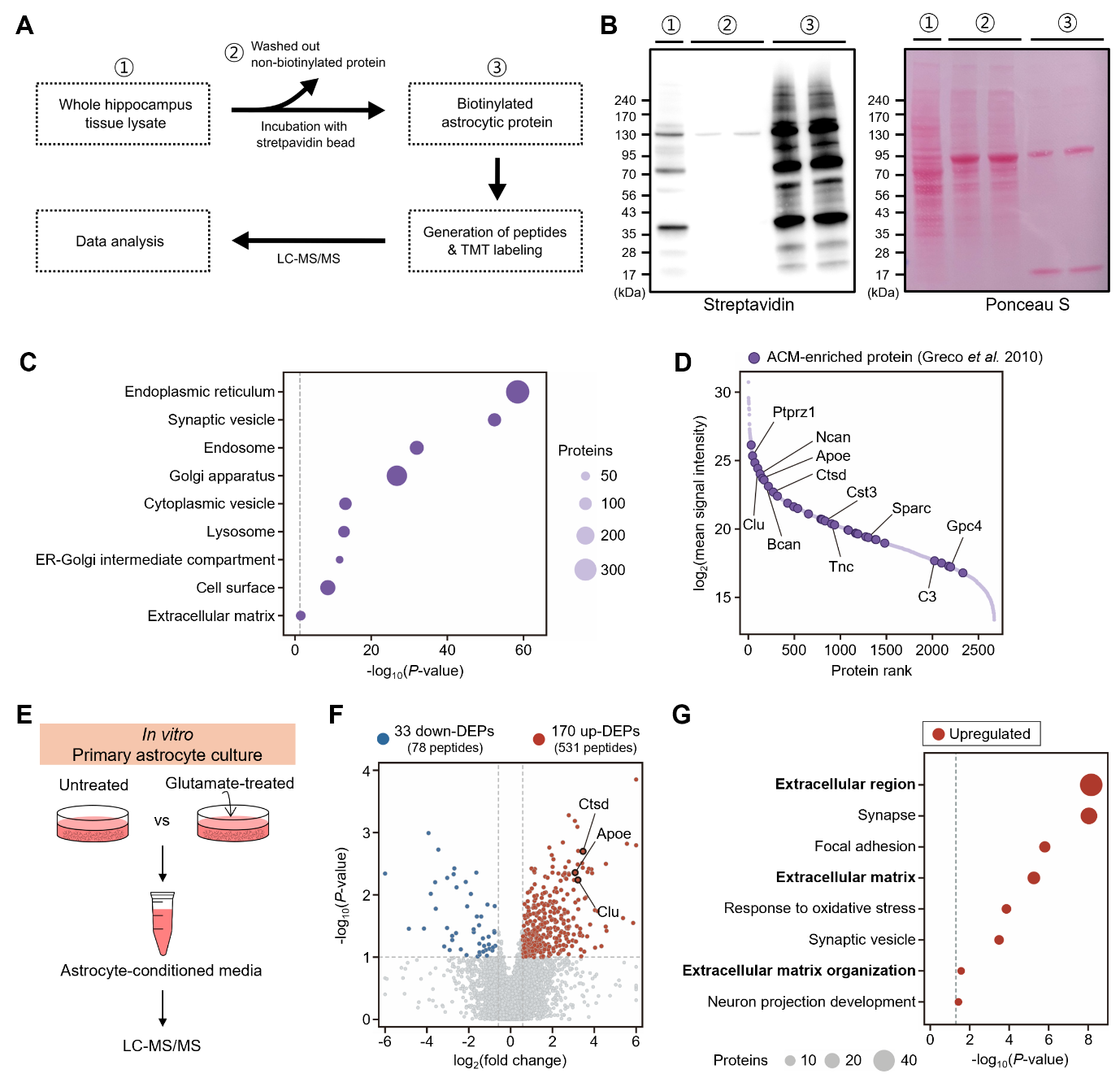
**Figure S3. Identification of learning-induced astrocytic proteins.**

**A**. Experimental scheme showing the process of astrocytic protein identification. **B**. Representative western blot and Ponceau S staining images showing purification of biotinylated proteins by using streptavidin beads. Each number indicates the corresponding step in **A**. **C.** Representative gene ontology terms enriched in the hippocampal astrocytic proteome. Circle size indicates the number of proteins. The dashed line indicates *P* = 0.05. **D.** Rank–abundance plot of the hippocampal astrocytic proteome. Proteins are ranked by mean signal intensity. ACM-enriched proteins^24^ are shown in purple. **E.** Schematic illustration of the workflow for identifying astrocyte-secreted proteins *in vitro* using primary ACM. **F.** Volcano plot of peptides quantified in the ACM of glutamate-treated versus untreated astrocytes. The x- and y-axes present the log_2_-fold change and –log_10_(*P-*value), respectively. Red and blue dots represent upregulated and downregulated proteins, respectively; gray dots represent proteins without significant differences in expression. Representative ECM-related proteins are highlighted. **G.** Representative gene ontology terms enriched in upregulated DEPs of the ACM secretome. Circle size indicates the number of proteins. The dashed line indicates *P* = 0.05.

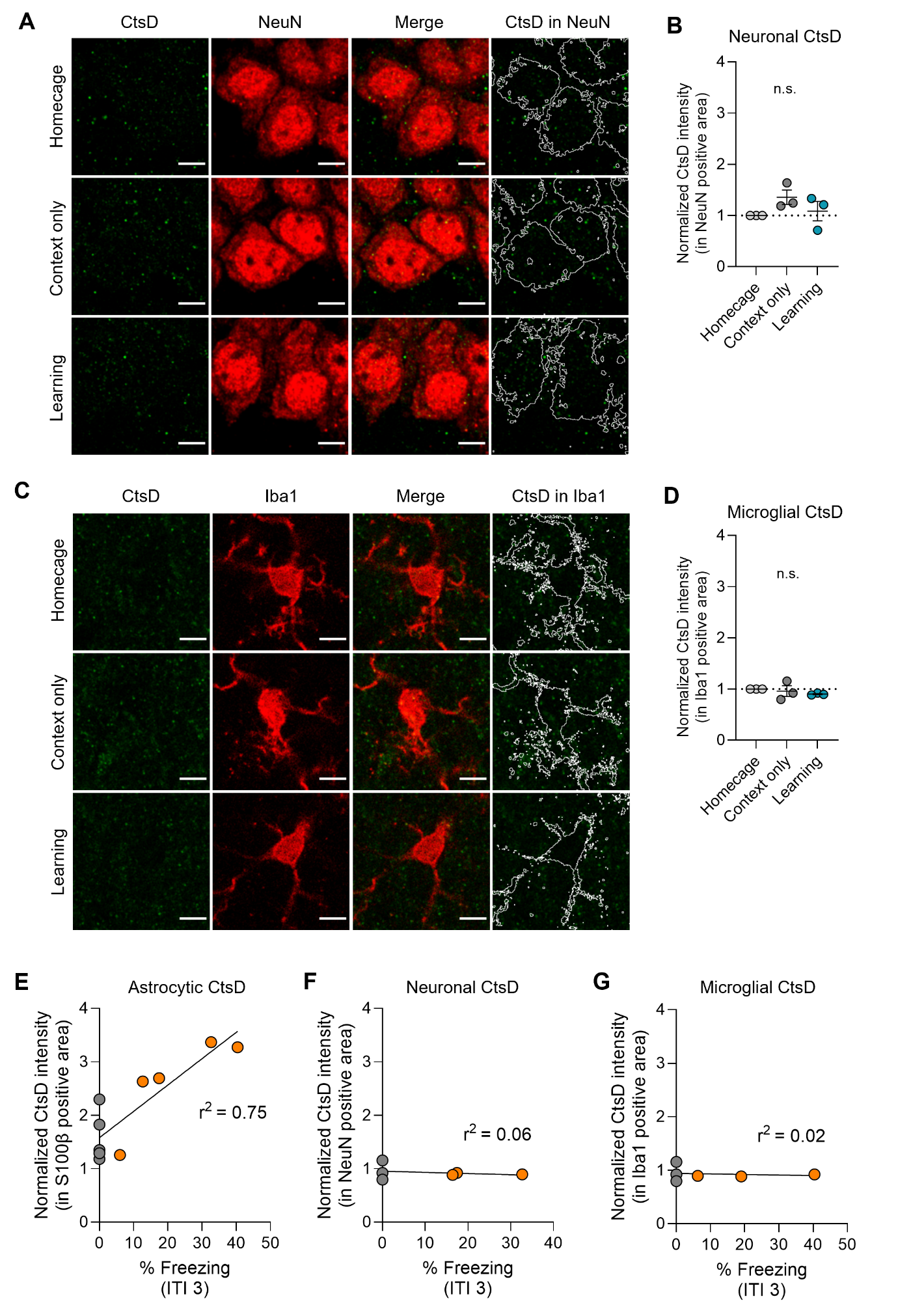
 **Figure S4. Astrocyte-selective elevation of CtsD following learning.**

**A.** Representative confocal images of the hippocampal CA1 region stained for CtsD (green) and a neuronal marker (NeuN; red). White outlines indicate NeuN-positive area. Scale bar, 5 μm. **B.** Quantification of CtsD fluorescence intensity within neuron areas. Each data point represents the mean value from one mouse. One-way ANOVA followed by Tukey’s multiple-comparisons test. Homecage, n = 3 mice; Context only, n = 3 mice; Learning, n = 3 mice. **C.** Representative confocal images of the hippocampal CA1 region stained for CtsD (green) and a microglial marker (Iba1; red). White outlines indicate microglia area. Scale bar, 5 μm. **D.** Quantification of CtsD fluorescence intensity within microglia areas. Each data point represents the mean value from one mouse. One-way ANOVA followed by Tukey’s multiple-comparisons test. Homecage, n = 3 mice; Context only, n = 3 mice; Learning, n = 3 mice. **E–G.** Correlation between freezing percentage during the third intertrial interval (ITI 3) and normalized CtsD intensity within S100β-positive (**e**), NeuN-positive (**f**), and Iba1-positive (**g**) areas. Lines indicate linear regression fits; r² values are shown. Each dot represents an individual mouse.

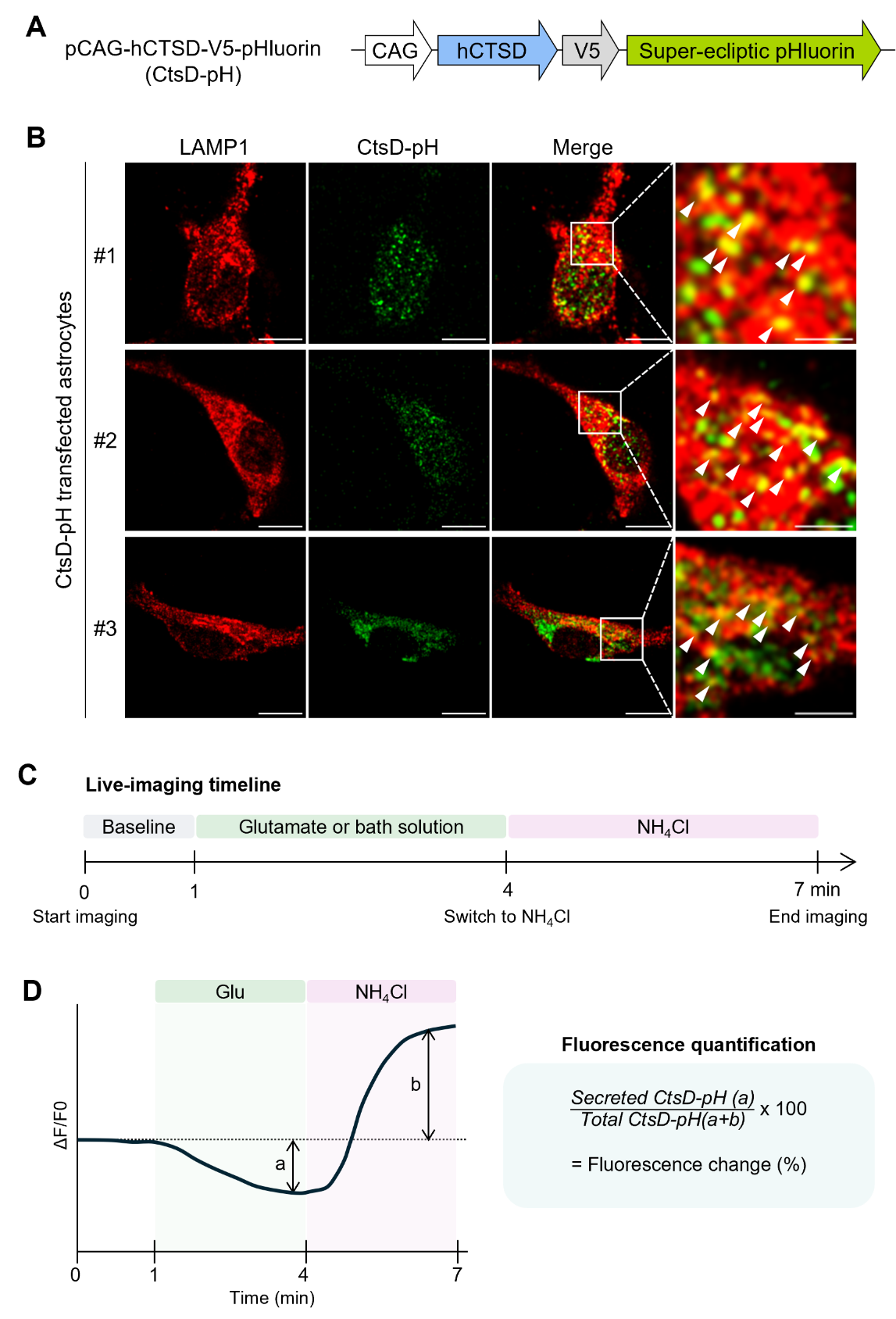

**Figure S5. CtsD-pH construct design, subcellular localization, and live-cell imaging in astrocytes.**

**A.** Illustration of the pCAG-hCTSD-V5-pHluorin construct (CtsD-pH). **B.** Representative confocal images of CtsD-pH-transfected astrocytes showing colocalization of CtsD-pH with the lysosomal marker (LAMP1). **C.** Timeline for CtsD-pH live-cell imaging. **D.** Schematic diagram explaining how to calculate normalized CtsD-pH fluorescence changes (fluorescence change %).

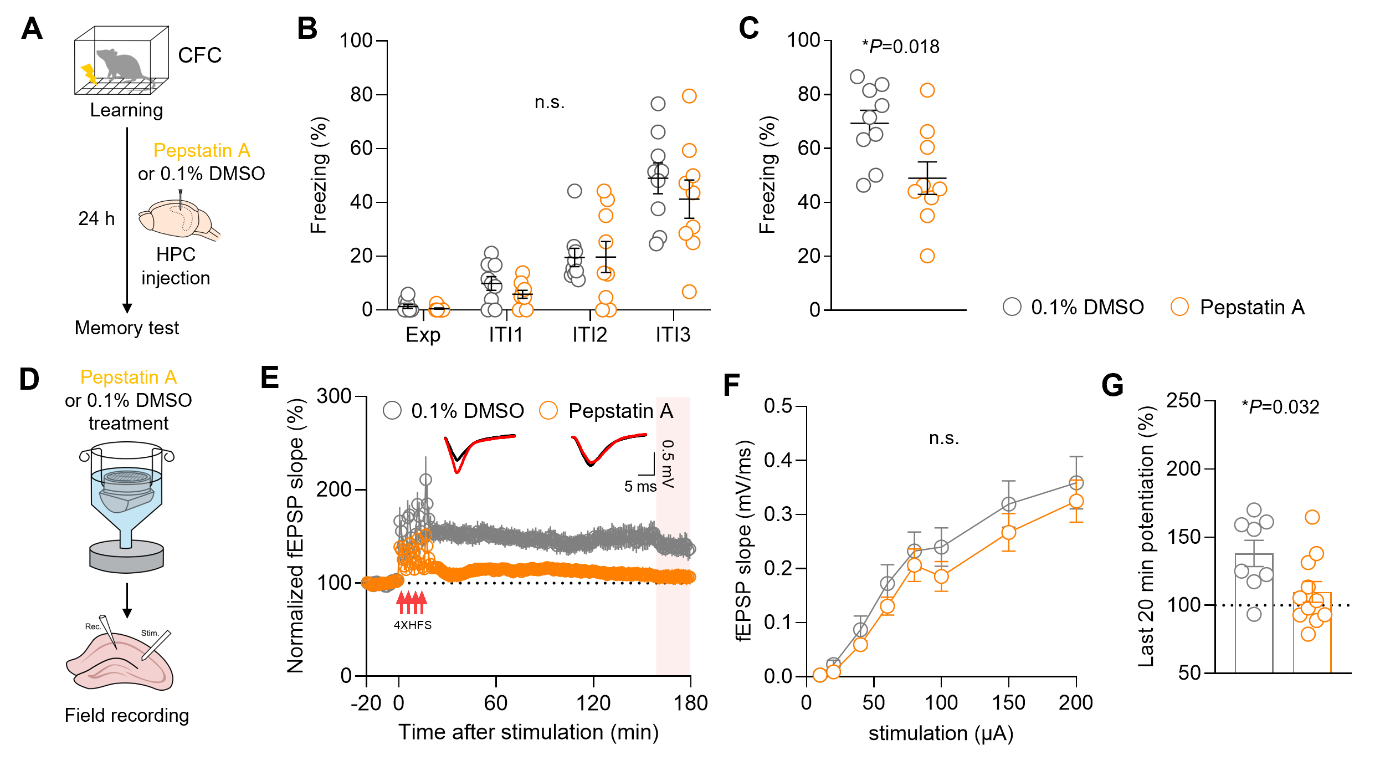

**Figure S6. Pepstatin A-mediated inhibition of extracellular CtsD impairs contextual fear memory and hippocampal LTP.**

**A.** Experimental scheme for CFC and the timing of pepstatin A injection. **B.** Percentage of time spent freezing during CFC training. Exp, exploration; ITI, intertrial interval. Two-way ANOVA followed by Tukey’s multiple-comparisons test. **C.** Percentage of time spent freezing during the memory test of CFC. Unpaired two-tailed Student’s t test. **B, C.** 0.1% DMSO, n = 9 mice; pepstatin A, n = 9 mice. **D.** Schematic diagram of *ex vivo* treatment of wild-type hippocampal slices with 0.1% DMSO or pepstatin A. **E.** Time courses of fEPSP responses before and after HFS (red arrows) in hippocampal slices treated with 0.1% DMSO or pepstatin A. Representative traces obtained during baseline recording (black) and 160–180 min after HFS (red) are shown. **F.** Input–output relationships of fEPSP slope in hippocampal slices treated with 0.1% DMSO or pepstatin A. Two-way ANOVA followed by Tukey’s multiple-comparisons test. **G.** Quantification of fEPSP potentiation during 160–180 min after HFS (pink box in **E**). Unpaired two-tailed Student’s *t* test. **E–G.** 0.1% DMSO, n = 8 slices from 4 mice; pepstatin A, n = 12 slices from 3 mice.

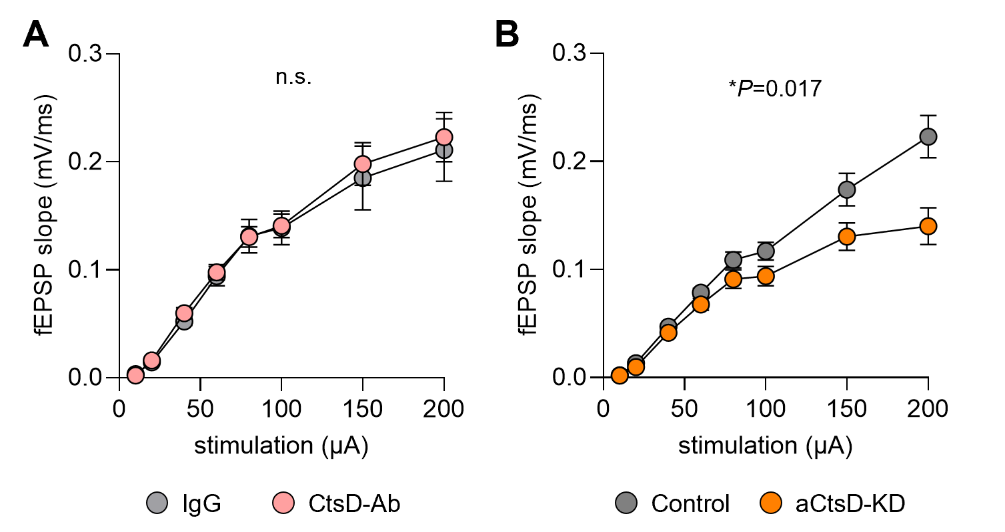

**Figure S7. Astrocytic CtsD knockdown, but not acute CtsD inhibition, impairs hippocampal synaptic transmission.**

**A.** Input–output relationships of fEPSP slope in hippocampal slices treated with control IgG or CtsD-Ab. Two-way ANOVA followed by Tukey’s multiple-comparisons test. IgG, n = 12 slices from 4 mice; CtsD-Ab, n = 11 slices from 5 mice. **B.** Input–output relationships of fEPSP slope in hippocampal slices from Control or aCtsD-KD mice. Two-way ANOVA followed by Tukey’s multiple-comparisons test. Control, n = 16 slices from 5 mice; aCtsD-KD, n = 14 slices from 5 mice.

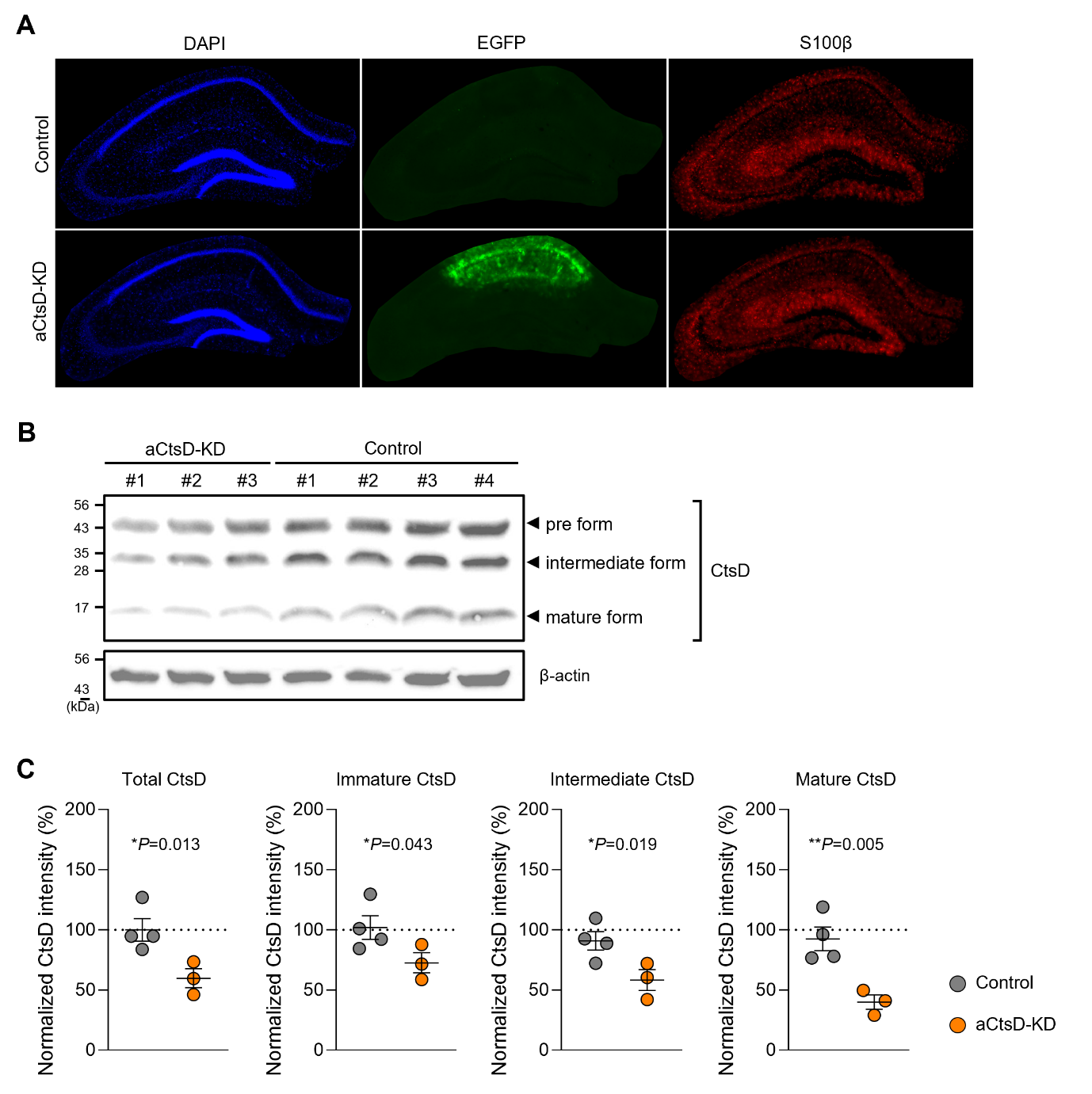

**Figure S8. Validation of astrocyte-targeted CtsD knockdown in the hippocampus.**

**A.** Representative fluorescence images of the hippocampus showing DAPI (blue), EGFP (green), and S100β (red) in Control and aCtsD-KD mice. **B.** Representative western blots showing precursor, intermediate, and mature forms of CtsD in hippocampal lysates from Control and aCtsD-KD mice. β-actin was used as a loading control. **C.** Quantification of total CtsD and its precursor, intermediate, and mature forms. Unpaired two-tailed Student’s t test. Control, n = 4 mice; aCtsD-KD, n = 3 mice.

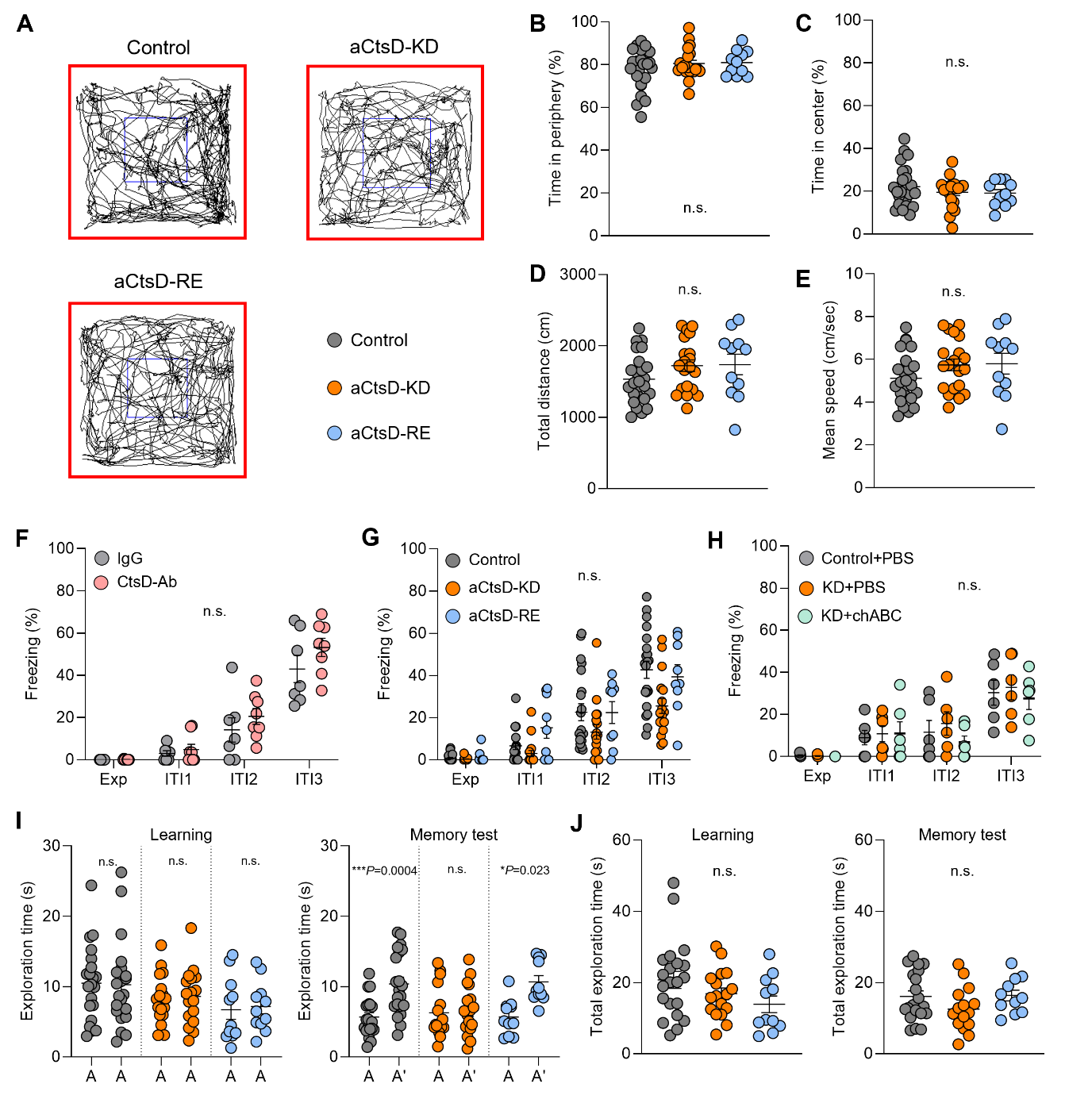
 **Figure S9. Astrocytic CtsD manipulation does not alter locomotor activity, anxiety-like behavior, or baseline learning performance.**

**A.** Representative movement trajectories of Control, aCtsD-KD, and aCtsD-RE mice in the open-field test. **B, C.** Percentage of time spent in the periphery (**B**) and center (**C**) of the open-field arena. **D, E.** Total distance traveled (**D**) and mean speed (**E**) during the open-field test. **B–E.** One-way ANOVA followed by Tukey’s multiple-comparisons test. Control, n = 25 mice; aCtsD-KD, n = 20 mice; aCtsD-RE, n = 11 mice. **F–H.** Percentage of time spent freezing during CFC training. Two-way ANOVA followed by Tukey’s multiple-comparisons test. **F.** IgG, n = 7 mice; CtsD-Ab, n = 8 mice. **G.** Control, n = 21 mice; aCtsD-KD, n = 17 mice; aCtsD-RE, n = 9 mice. **H.** Control + PBS, n = 6 mice; aCtsD-KD + PBS, n = 6 mice; aCtsD-KD + ChABC, n = 6 mice. **I.** Time spent exploring each object during the learning and memory test phases of the NOLT in Control, aCtsD-KD, and aCtsD-RE mice. **J.** Total object exploration time during the learning and memory test phases of the NOLT. **I, J.** One-way ANOVA followed by Tukey’s multiple-comparisons test. Control, n = 21 mice; aCtsD-KD, n = 17 mice; aCtsD-RE, n = 11 mice.

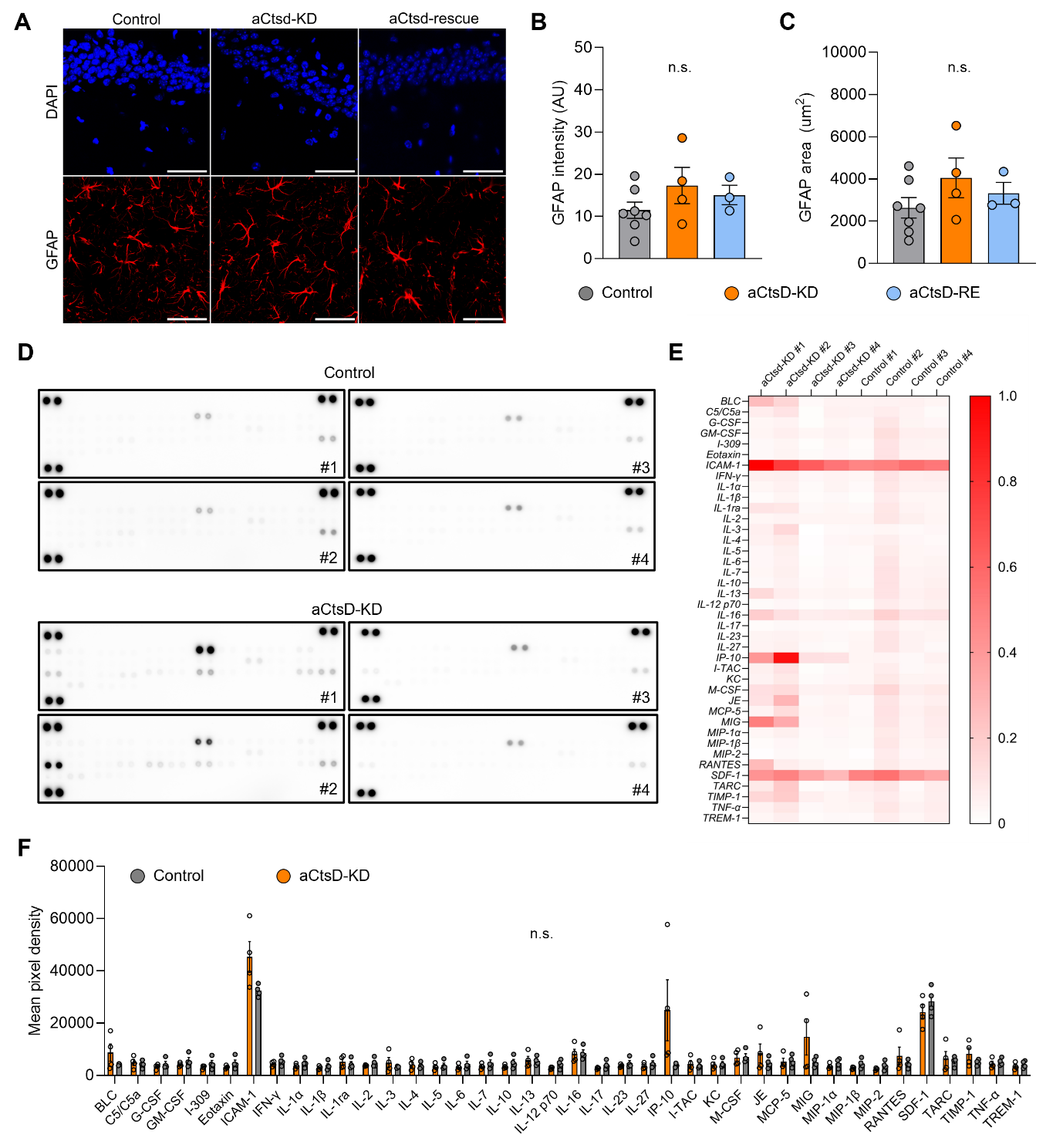

**Figure S10. Astrocytic CtsD knockdown does not trigger astrocyte reactivity or neuroinflammatory responses.**

**A.** Representative confocal images of the hippocampus stained for DAPI (blue) and GFAP (red) in Control, aCtsD-KD, and aCtsD-RE mice. Scale bar, 50 μm. **B, C.** Quantification of GFAP fluorescence intensity (**B**) and GFAP-positive area (**C**). One-way ANOVA followed by Tukey’s multiple-comparisons test. Control, n = 6 mice; aCtsD-KD, n = 4 mice; aCtsD-RE, n = 3 mice. **D.** Cytokine and chemokine array images of hippocampal lysates from Control and aCtsD-KD mice. **E.** Heatmap showing min–max-normalized cytokine and chemokine array signal intensities. **F.** Quantification of cytokine and chemokine array signals expressed as mean pixel density. Multiple unpaired t tests with FDR control using the two-stage step-up method of Benjamini, Krieger and Yekutieli (Q = 1%). Control, n = 4 mice; aCtsD-KD, n = 4 mice.

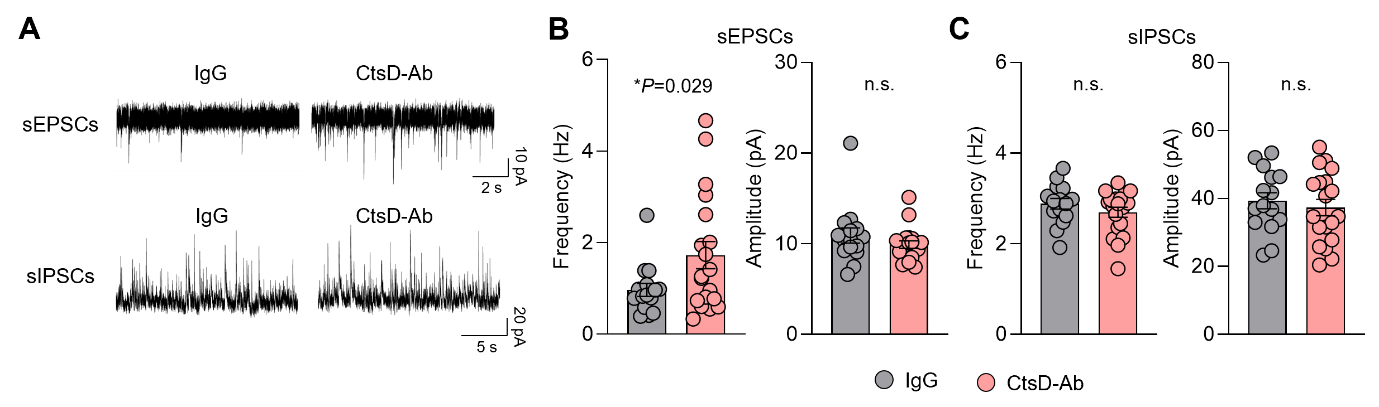

**Figure S11. Acute extracellular CtsD neutralization selectively increases sEPSCs.**

**A.** Representative traces of sEPSCs (top) and sIPSCs (bottom) in IgG- or CtsD-Ab-treated hippocampal slices prepared from wild-type mice. **B, C.** Quantification of the frequency and amplitude of sEPSCs (b) and sIPSCs (c). Unpaired two-tailed Student’s *t* test. IgG, n = 15 cells from 3 mice; CtsD-Ab, n = 19 cells from 4 mice.

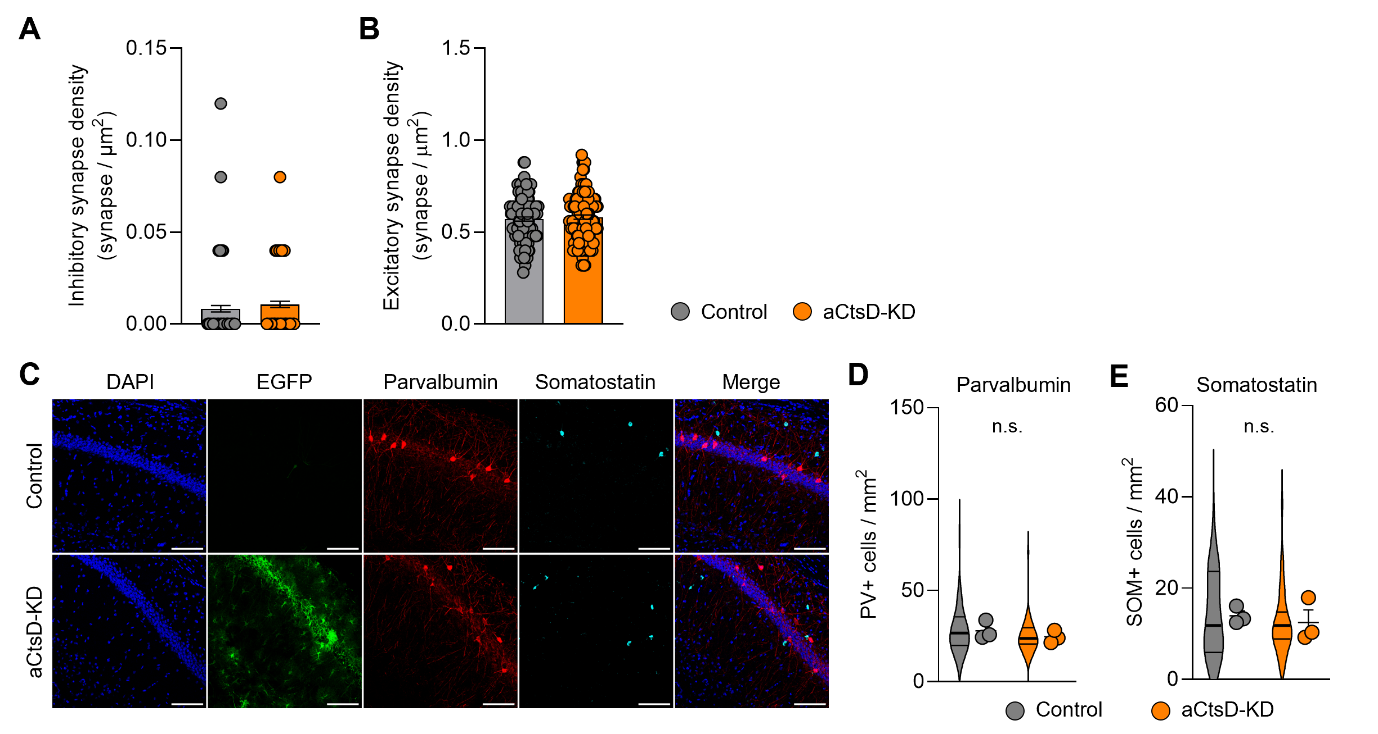
 **Figure S12. Ultrastructural analysis reveals unchanged excitatory and inhibitory synapse densities in aCtsD-KD mice.**

**A, B**. Quantification of inhibitory (a) and excitatory(b) synapses using TEM images. Unpaired two-tailed Student’s *t* test. Control, n = 120 images from 4 mice; aCtsD-KD, n = 120 images from 4 mice. **C.** Representative confocal images of the hippocampal CA1 region showing DAPI (blue), EGFP (green), PV (red), and SOM (cyan) in Control and aCtsD-KD mice. Scale bar, 100 μm. **D, E.** Quantification of PV-positive (b) and SOM-positive (c) cell density in the hippocampal CA1 region. Unpaired two-tailed Student’s *t* test. Control, n = 3 mice (46 images); aCtsD-KD, n = 3 mice (47 images).

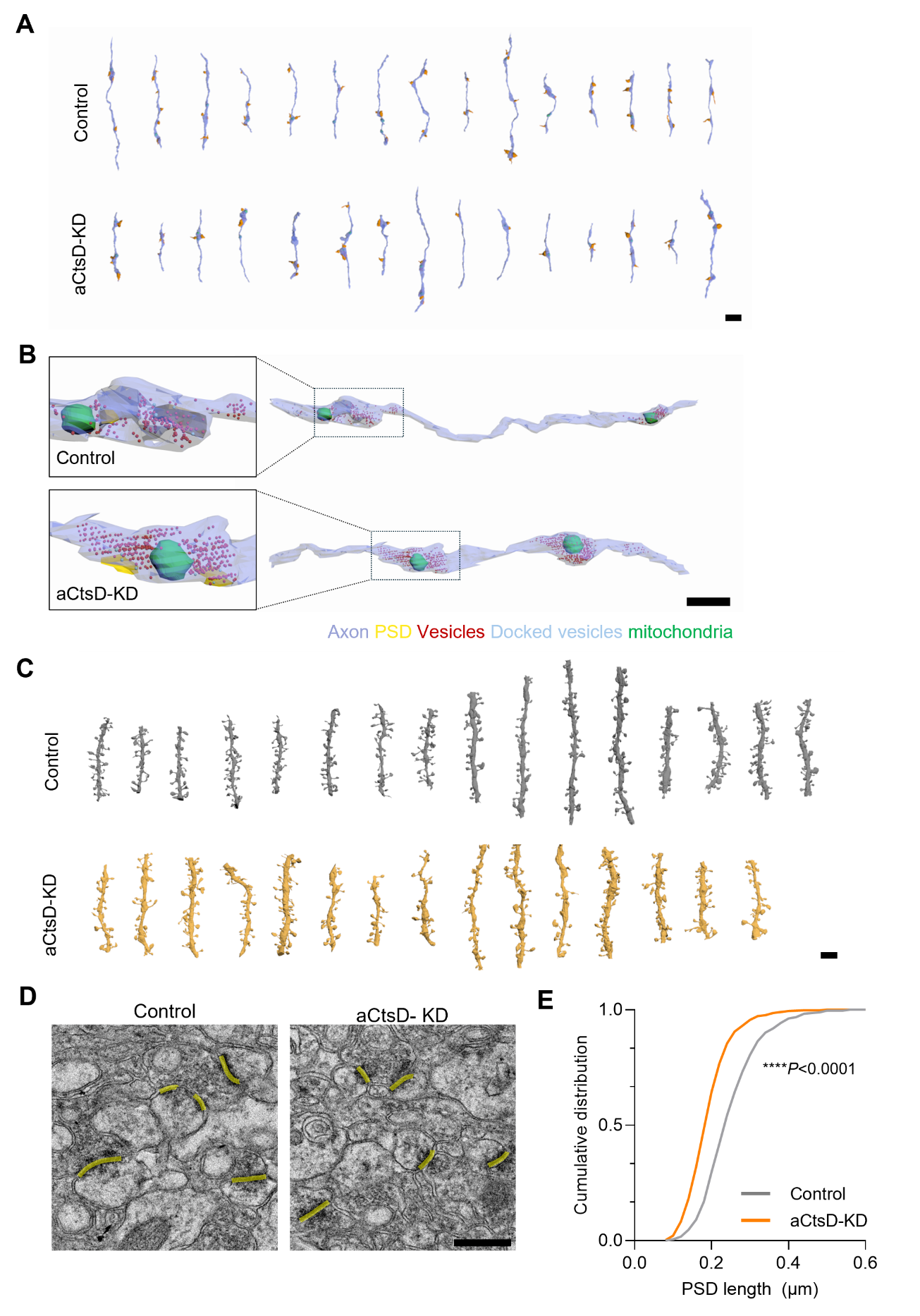
**Figure S13. Ultrastructural analysis of hippocampal synapses in aCtsD-KD mice.**

**A.** Representative 3D reconstructions of axons and their contacting postsynaptic spines in the vicinity of astrocytes in the CA1 region of Control and aCtsD-KD mice. Scale bar, 2 μm. **B.** Representative 3D reconstructions of excitatory presynaptic boutons from Control and aCtsD-KD mice, showing the spatial organization of synaptic vesicles (red), docked vesicles (light blue), PSDs (yellow), and mitochondria (green). Scale bar = 1 μm. **C.** Representative 3D reconstructions showing dendritic spine morphology of CA1 pyramidal neurons in Control and aCtsD-KD mice. Scale bar = 2 μm. **D.** Representative transmission electron microscopy images showing postsynaptic density (PSD) lengths, indicated by yellow lines. Scale bar, 0.5 μm. **E.** Cumulative distribution of PSD lengths. Kolmogorov-Smirnov test. Control, n = 507 PSDs from 4 mice; aCtsD-KD, n = 507 PSDs from 4 mice.

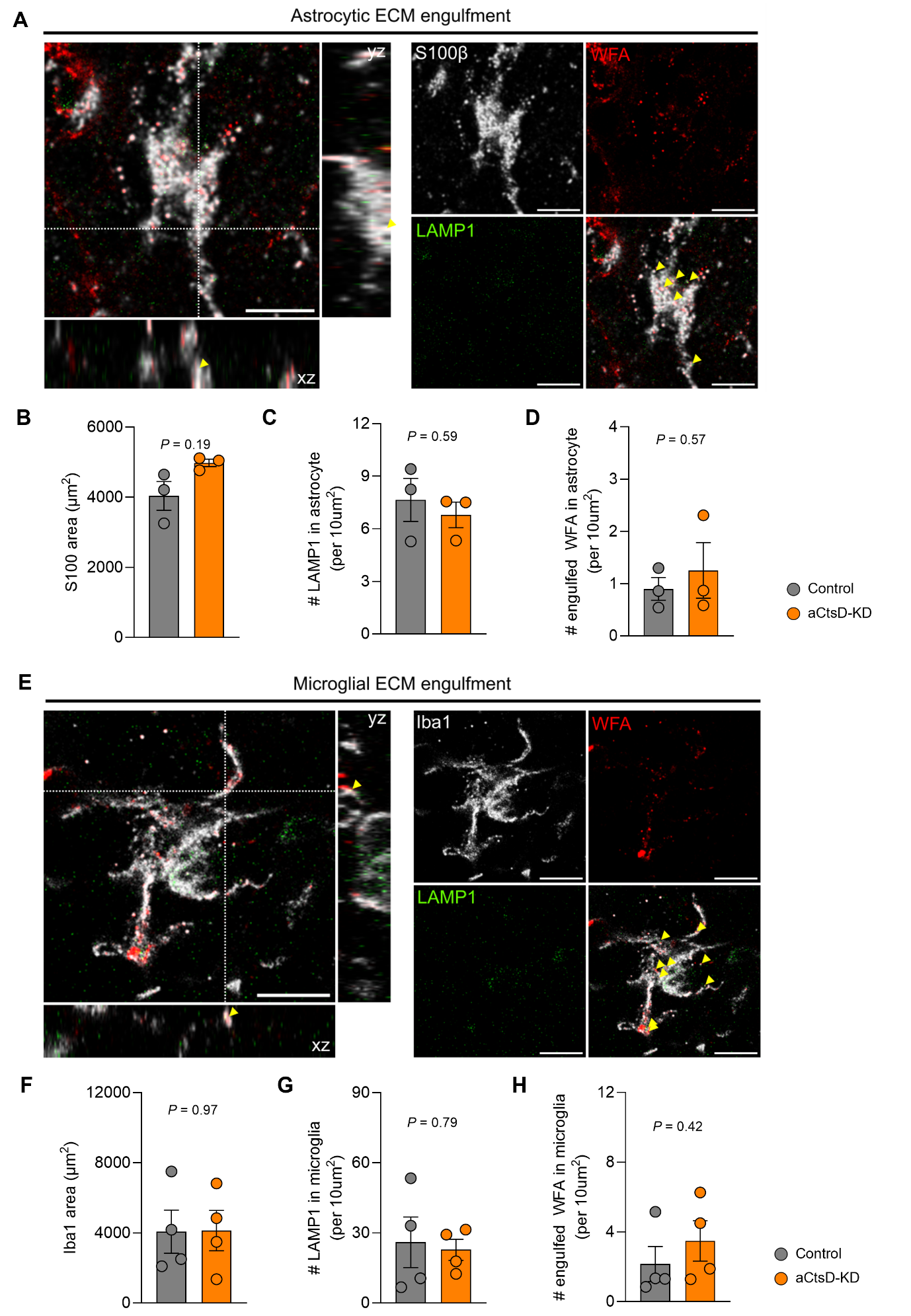

**Figure S14. Astrocytic CtsD knockdown does not alter microglial or astrocytic ECM engulfment.**

**A.** Representative confocal images showing S100β (white), WFA (red), and LAMP1 (green), with orthogonal views. Yellow arrowheads indicate engulfed WFA-positive puncta. Scale bars, 10 μm. **B–D.** Quantification of S100β-positive area (**B**), LAMP1-positive puncta within astrocytes (**c**), and engulfed WFA-positive puncta within astrocytes (**D**). Unpaired two-tailed Student’s *t* test. Control, n = 3 mice; aCtsD-KD, n = 3 mice. **e.** Representative confocal images showing Iba1 (white), WFA (red), and LAMP1 (green), with orthogonal views. Yellow arrowheads indicate engulfed WFA-positive puncta. Scale bars, 10 μm. **F–H.** Quantification of Iba1-positive area (**F**), LAMP1-positive puncta within microglia (**g**), and engulfed WFA-positive puncta within microglia (**H**). Unpaired two-tailed Student’s *t* test. Control, n = 3 mice; aCtsD-KD, n = 3 mice.

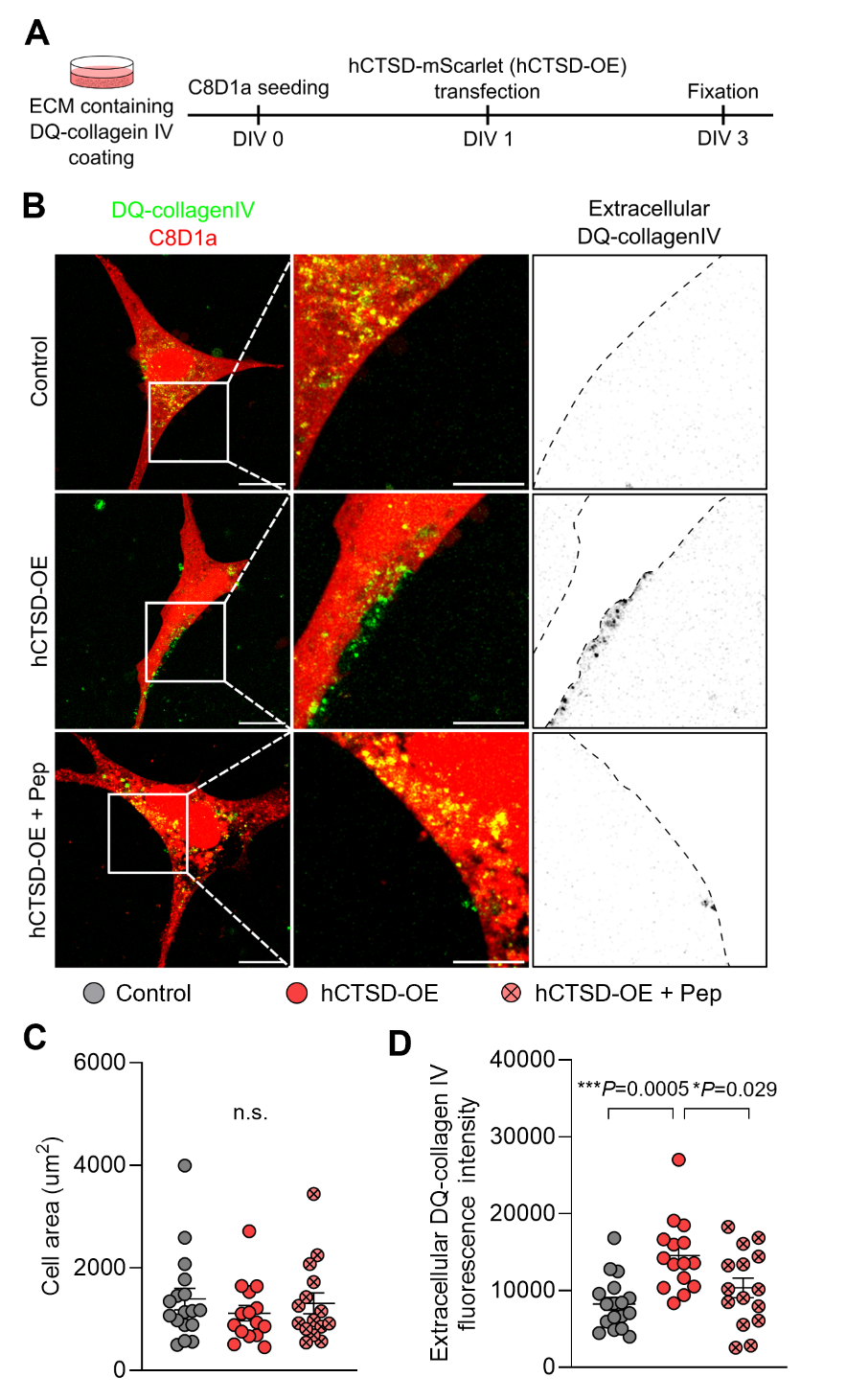

**Figure S15. hCTSD overexpression promotes extracellular degradation of collagen IV in astrocyte cultures.**

**A.** Experimental scheme for assessing extracellular collagen IV degradation following hCTSD overexpression. **B.** Representative fluorescence images of Control or hCTSD-overexpressing (hCTSD-OE) astrocytes in the absence or presence of pepstatin A. DQ-collagen IV fluorescence is shown in green, and astrocytes are shown in red. **c, d.** Quantification of cell area (**C**) and extracellular DQ-collagen IV integrated fluorescence intensity (**D**). One-way ANOVA followed by Tukey’s multiple-comparisons test. Control, n = 17 cells; hCTSD-OE, n = 15 cells; hCTSD-OE + Pep, n = 15 cells from 3 independent experiments.

**Table 5. Electrophysiological properties of the recorded neurons**

|  | Membrane capacitance (pF) | Membrane resistance (mΩ) | Holding current  (pA) |
| --- | --- | --- | --- |
| Control | 35.73 ± 2.147 | 99.74 ± 5.383 | -19.55 ± 6.446 |
| aCtsD-KD | 38.55 ± 2.575 | 104.2 ± 5.896 | -17.75 ± 6.720 |
| *P* value | *P* = 0.4146 | *P* = 0.5838 | *P* = 0.8492 |

Unpaired two-tailed Student’s *t* test. Mean ± SEM.

- Internal solution: CsMeSO_3_-based solution

|  | Membrane capacitance (pF) | Membrane resistance (mΩ) | Holding current  (pA) |
| --- | --- | --- | --- |
| Control | 28.10 ± 1.263 | 124.8 ± 7.593 | -37.14 ± 10.87 |
| aCtsD-KD | 24.64 ± 1.911 | 128.6 ± 5.197 | -41.34 ± 7.328 |
| *P* value | *P* = 0.1362 | *P* = 0.6814 | *P* = 0.7529 |

- Internal solution: CsCl-based solution

|  | Membrane capacitance (pF) | Membrane resistance (mΩ) | Holding current  (pA) |
| --- | --- | --- | --- |
| IgG | 52.86 ± 2.615 | 103.2 ± 5.184 | -29.40 ± 6.404 |
| CtsD-Ab | 51.61 ± 1.963 | 107.8 ± 4.179 | -24.34 ± 6.137 |
| *P* value | *P* = 0.7007 | *P* = 0.4915 | *P* = 0.5760 |

- Internal solution: CsMeSO_3_-based solution

**Table 6. Antibodies and reagent resources**

| Antibody / Reagent | Supplier | Catalog no. | Application | Dilution |
| --- | --- | --- | --- | --- |
| Mouse anti-V5 | Thermo fisher scientific | R960-25 | IHC, WB | 1:1000, 1:5000 |
| Rabbit anti-β-Actin-HRP | Cell Signaling | 5125S | WB | 1:4000 |
| Streptavidin-HRP | Thermo fisher scientific | 21124 | WB | 1:10000 |
| Streptavidin – 488 | Thermo fisher scientific | 32354 | ICC | 1:500 |
| Streptavidin – 555 | Thermo fisher scientific | 32355 | IHC | 1:500 |
| Streptavidin – 680 | Jackson ImmunoResearch | 016-620-084 | ICC | 1:200 |
| Rabbit anti-NeuN | Abcam | ab177487 | IHC | 1:1000 |
| Rabbit anti-Iba1 | FUJIFILM Wako | 019-19741 | IHC | 1:500 |
| Rabbit anti-S100 | DAKO | Z0311 | IHC | 1:500 |
| Guinea pig anti-S100β | Synaptic Systems | 287 004 | IHC | 1:500 |
| Mouse anti-Cathepsin D (C-5) | Santa Cruz Biotechnology | sc-377124 | IHC | 1:25 |
| Rabbit anti-Cathepsin D (E5V4H) | Cell Signaling | 74089S | WB | 1:1000 |
| Chicken anti-GFAP | MilliporeSigma | AB5541 | IHC | 1:500 |
| Chicken anti-Parvalbumin | Synaptic Systems | 195 006 | IHC | 1:500 |
| Rabbit anti-Somatostatin | ImmunoStar | 20067 | IHC | 1:500 |
| Rabbit anti-LAMP1 | Abcam | Ab24170 | IHC | 1:200 |
| Chicken anti-Iba1 | Synaptic Systems | 234 009 | IHC | 1:500 |
| Wisteria Floribunda Lectin (WFA, WFL), biotinylated | Vector laboratories | B-1355-2 | IHC | 1:200 |
| Guinea pig anti-VGAT | Synaptic Systems | 131 004 | IHC | 1:500 |
| Rabbit-GABA-A receptor α1 | Synaptic Systems | 224 203 | IHC | 1:500 |
| Rabbit-GABA-A receptor α2 | Synaptic Systems | 224 103 | IHC | 1:500 |
| Goat anti-rabbit alexa 405 | Invitrogen | A-31556 | IHC | 1:200 |
| Goat anti-rabbit alexa 488 | Invitrogen | A-11034 | IHC | 1:200 |
| Goat anti-rabbit alexa 568 | Invitrogen | A-11011 | IHC | 1:200 |
| Goat anti-rabbit alexa 647 | Invitrogen | A-21245 | IHC | 1:200 |
| Goat anti-chicken alexa 647 | Invitrogen | A-21449 | IHC | 1:200 |
| Goat anti-guinea pig alexa 546 | Invitrogen | A-11074 | IHC | 1:200 |
| Anti-mouse-IgG-HRP | Bio-Rad | 170-6546 | WB | 1:10000 |
| Anti-rabbit-IgG-HRP | Bio-Rad | 170-6515 | WB | 1:4000 |
| Normal goat serum | Jackson ImmunoResearch | 005–000-121 |  |  |
| Antifade Mounting medium with DAPI | Vector Laboratories | H-1200 |  |  |
| Antifade Mounting medium | Vector Laboratories | H-1400 |  |  |
| DAPI | Sigma-Aldrich | D9542 |  |  |
| Skim milk powder | Sigma-Aldrich | 70166 |  |  |
| Bovine Serum Albumin | Sigma-Aldrich | A7906 |  |  |
| SuperSignal West Pico PLUS | Thermo fisher scientific | 34580 |  |  |
| EZ-Western Lumi Femto | DoGen | DG-WF200 |  |  |
| Tween 20 | Sigma-Aldrich | P9416 |  |  |
| TritonX-100 | Sigma-Aldrich | X100 |  |  |
| 4x Laemmli sample buffer | Bio-Rad | 1610747 |  |  |
| Dimethyl sulfoxide (DMSO) | Sigma‒Aldrich | D8418 |  |  |
| Ponceau S solution | ROCKLAND | MB-072-0500 |  |  |

- ICC, Immunocytochemistry; IHC, Immunohistochemistry; WB, Western blot; HRP, Horseradish Peroxidase
